# Chromosomal inversions harbour excess mutational load in the coral, *Acropora kenti,* on the Great Barrier Reef

**DOI:** 10.1101/2024.02.19.580031

**Authors:** Jia Zhang, Nadja M. Schneller, Matt A. Field, Cheong Xin Chan, David J. Miller, Jan M. Strugnell, Cynthia Riginos, Line Bay, Ira Cooke

## Abstract

The future survival of coral reefs in the Anthropocene depends on the capacity of corals to adapt as oceans warm and extreme weather events become more frequent. Targeted interventions designed to assist evolutionary processes in corals require a comprehensive understanding of the distribution and structure of standing variation, however, efforts to map genomic variation in corals have so far focussed almost exclusively on SNPs, overlooking structural variants that have been shown to drive adaptive processes in other taxa. Here we show that the reef-building coral, *Acropora kenti* (syn. tenuis) harbors at least five large, highly polymorphic structural variants, all of which exhibit signatures of strongly suppressed recombination in heterokaryotypes, a feature commonly associated with chromosomal inversions.

Based on their high minor allele frequency, uniform distribution across habitats, and elevated genetic load, we propose that these inversions in *A. kenti* are likely to be under balancing selection. An excess of SNPs with high impact on protein coding genes within these loci elevates their importance both as potential targets for adaptive selection and as contributors to genetic decline if coral populations become fragmented or inbred in future.

## Introduction

Coral reefs are hyperdiverse marine ecosystems that provide crucial ecosystem services to millions of people throughout the tropics. Threats to coral reefs, particularly from ocean warming have placed the evolutionary biology of corals into the spotlight because it is now clear that their long-term survival depends on whether they can adapt to keep pace with climate change (*1–4*). Decision making in relation to protected area design(*5*), genetic interventions(*6*) and reef restoration(*1*) must therefore be informed by a sound understanding of the factors that shape genetic diversity in corals(*7*).

Over the past two decades, adoption of population genomic approaches has greatly improved our understanding of evolutionary processes in corals. Using SNP and microsatellite markers many studies have identified instances of fine-scale population structure and cryptic speciation (*8–13*). More recently, the adoption of dense SNP marker sets and whole genome sequencing has started to reveal the origins and drivers of divergence(*14*, *15*) as well as the architecture of key traits such as heat tolerance(*16*). So far, however, all population genomic work in corals has relied exclusively on SNPs, ignoring structural variants, such as inversions. Inversions are a particularly important form of structural variation because they can be large (multiple Mb in genomic extent) and can strongly suppress recombination between inverted and ancestral karyotypes(*17*). These characteristics make them potent evolutionary modifiers that can facilitate local or clinal adaptive processes(*18–20*), often have strong phenotypic effects(*21*, *22*), and may capture a large fraction of the standing genetic variation in some species (*23*, *24*).

The fact that inversions inhibit recombination over large genomic regions provides a mechanism for local adaptation under gene flow(*25*) because an inversion that captures a locally favourable combination of alleles will be protected from recombination with less favourable alleles on the alternate arrangement(*26*, *27*). Initially predicted from simple theoretical models(*27*), this idea is now supported by an increasing number of examples across a wide variety of taxa(*18*, *22*, *28*, *29*) in which alternate arrangements of an inversion are found to diverge in frequency between ecotypes. Inversion polymorphisms that become established via such spatially divergent selection (classified as type I(*30*)) have received considerable attention in the literature, perhaps because they can easily be detected as large blocks of sharply elevated F_ST_ between ecotypes(*31*). Recent theoretical work also suggests that such type I inversions are more likely to be large(*32*) and therefore more discoverable than those established via other mechanisms.

There is increasing recognition that balancing selection plays a crucial role in the establishment and persistence of many inversions(*18*, *19*, *30*, *33*). Several recent studies have shown that accumulation of recessive deleterious mutations within inversions can directly favour the hetero-karyotype (*22*, *33–35*), preventing inversions from reaching fixation and supporting long-term persistence at high frequency throughout the population. Since inversions maintained by this mechanism need not segregate strongly across ecotypes they may be more challenging to identify, and their ecological roles less obvious, however, evidence is emerging in support of the idea that inversions of this type (type II; (*30*)) may form an important reservoir of genetic variation over and above that of the collinear genome. Recent theoretical work(*35*) and empirical observations in sunflowers(*36*) suggest that type II inversions may accumulate mutational load at a higher rate than type I inversions because the relative lack of homokaryotypes inhibits purging of deleterious alleles. Since selective forces on inversions may change over time, the reservoir of variation accumulated within type II inversions may eventually become a target for positive selection. This idea is supported by a recent global analysis(*20*) of the In(3R)Payne inversion in *Drosophila melanogaster* which occurs as a balanced polymorphism in its ancestral African population but has now formed sharp latitudinal clines underlying climate adaptation in North America(*37*) and Australia(*38*).

Although large polymorphic inversions have now been observed across a wide variety of taxa(*23*, *24*, *39–41*), their role in corals, and more broadly in cnidarians, remains an open question. Evidence from comparative genomics has shown that several large inversions have accumulated between species of the genus *Acropora*(*42*), however, the prevalence of polymorphic inversions within cnidarian species is unknown, and might vary widely between species, as it does in *Drosophila* (*43*). There is growing consensus that some important traits such as bleaching tolerance in corals are likely to be controlled by many unlinked genes of small effect(*16*), however, inversions would complicate this paradigm because they can capture multiple loci that collectively have a large influence on adaptive and speciation processes(*19*). Since inversions can also harbour recessive deleterious variation it is possible that they could play a role in genetic decline, for example, if a population becomes fixed for an inversion after a bottleneck, exposing recessive phenotypes via homozygosity. A comprehensive understanding of the contribution of inversions and other structural variants to standing genetic variation in corals is therefore required in order to project their capacity to adapt to climate change, forecast the consequences of genetic decline, and to plan genomic interventions.

We set out to identify and characterise inversions and other genome-wide patterns of genetic variation, in the reef-building coral *Acropora kenti (syn. tenuis)*, sampled from five inshore and four offshore reefs along the same 500km section of the central Great Barrier Reef (GBR). *Acropora kenti*, previously classified as *Acropora tenuis*(*44*) has been the focus of genetic, developmental and physiological studies for many years (*12*, *45*, *46*) and is the target of research on potential genomic interventions to mitigate climate change impacts(*47*). Our study sites were chosen to include a contrast between inshore and offshore reefs which are spatially separated but likely to experience gene flow. Inshore reefs experience higher turbidity, more variable temperatures and much greater terrestrial influence (nutrients, pollution and freshwater runoff) than offshore(*48–50*). Evidence from previous studies suggests that there are also differences in dominant algal symbionts harboured by corals at different reefs in this region(*12*, *46*). While these environmental gradients provide selective pressures that might promote the formation of locally adapted ecotypes, they are also subject to high levels of gene flow which would oppose local adaptation. Our goal was to identify and characterise any polymorphic inversions present within this *A. kenti* population, determine their roles (if any) in promoting local adaptation and as contributors to standing genetic variation.

Using patterns of heterozygosity, linkage disequilibrium and local genetic structure we identified at least five inversions circulating at high frequency (MAF>0.17) and ranging in size from 0.2 to 2Mb. None of these five showed patterns of elevated differentiation between inshore/offshore or between colonies with different dominant symbionts that would demonstrate a role in local adaptation. Instead, our analyses show that all these inversions are highly polymorphic, harbour an excess of mutations with high impacts on protein coding genes, and are deficient in homokaryotypes. This combination of characteristics is most consistent with inversions that are maintained by forms of balancing selection that directly favour the heterokaryotypic state such as associative overdominance. Our results highlight the fact that structural variants such as inversions may be prevalent in coral populations and play a significant role in structuring standing genetic variation.

## Results

### Population structure and symbiont diversity

To facilitate detection and characterisation of inversions in *A. kenti* we used the ANGSD framework to call 3.8 million biallelic SNPs from shallow (2-5×) whole genome sequencing data across 208 genetically distinct colonies sampled from nine reefs in the central Great Barrier Reef (GBR) (Figure 1A). Full details of SNP calling and quality control are given in methods. Analyses of population structure and admixture with PCAngsd showed that eight of these reef populations formed a single genetic cluster that was distinct from Magnetic Island (Figure 1B). The strong distinction of Magnetic Island has been observed in previous whole genome sequencing (WGS) studies(*12*, *51*) and is thought to reflect divergence around 500kya-1Mya. Despite this divergence, we observed four highly admixed individuals, indicative of recent crosses between Magnetic Island and other reefs (Figure 1C). The eight non-Magnetic Island reefs in our study include one location (Pelorus Island) that overlaps with locations dominated by Cluster 1A identified by Matias et al(*12*), implying that this lineage occurs across the full length of the GBR, and can be found in both inshore and offshore locations(*12*).

**Figure 1:**
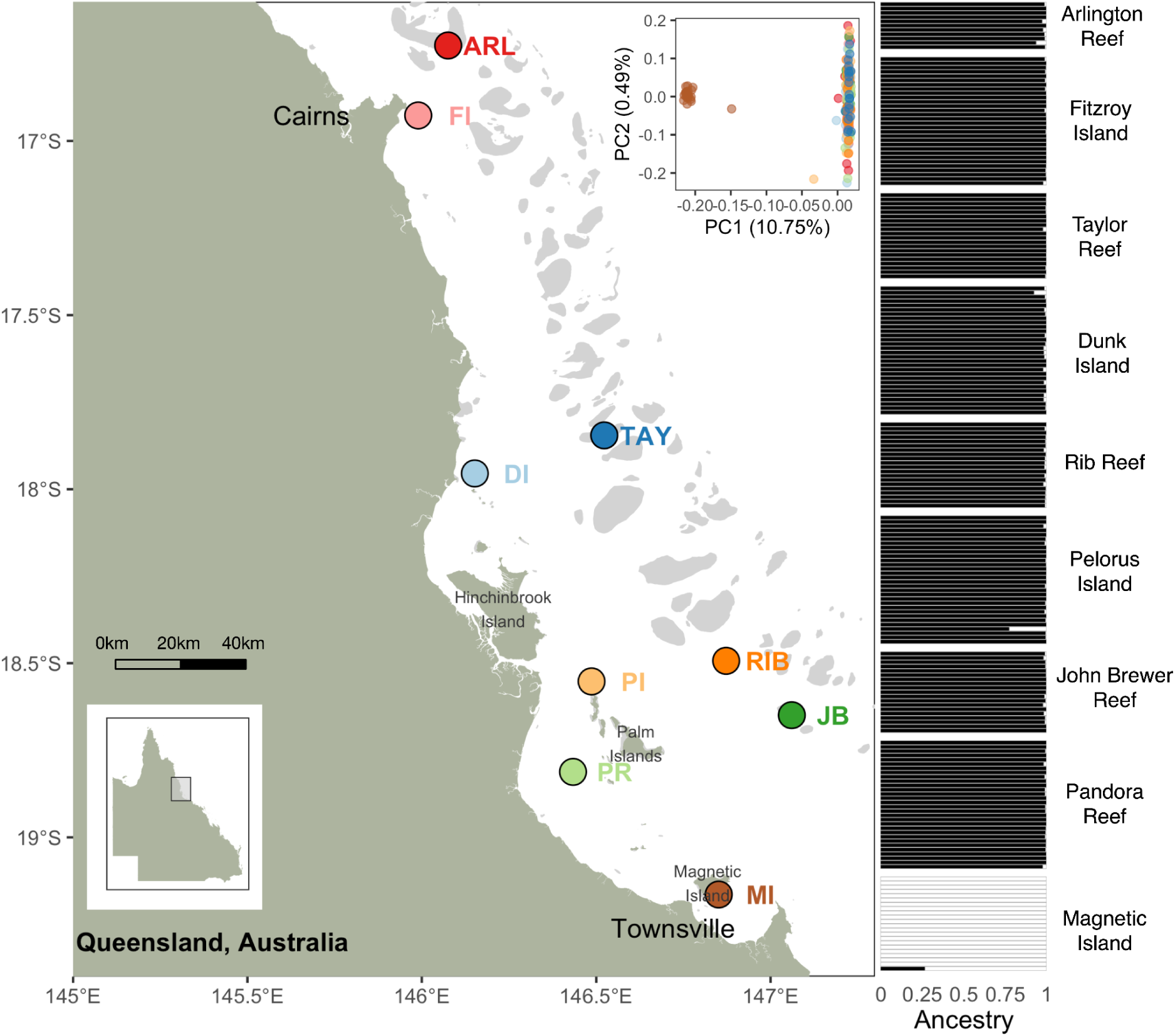
Population structure and sampling locations within the central Great Barrier Reef. Inset shows a Principal Components Analysis with two main clusters (Magnetic Island: left, Northern Reefs: right), for which the ancestry proportions are shown for each individual. Offshore reefs sequenced in this study are: Arlington Reef (ARL), Taylor Reef (TAY), Rib Reef (RIB), and John Brewer Reef (JB). Inshore reefs (sequenced in Cooke et al(*51*)) are: Fitzroy Island (FI), Dunk Island (DI), Palm Islands (PI), and Pandora reef (PR) and Magnetic Island (MI).

Among the eight non-Magnetic Island reefs, we found no clear evidence for genetic structure between reefs or between inshore vs offshore locations. This was evident in a PCA based on non-Magnetic Island data (Supplementary Figure S4) and a tree inferred from the identity by state (IBS) distance matrix between all pairs of samples (Supplementary Figure S5). Since inshore and offshore samples were sequenced in separate batches such lack of structure suggests that batch effects (if present) are likely to have minimal influence on population genetic inferences. Nevertheless, we found higher variability in individual heterozygosity estimates for inshore samples than for offshore (Supplementary Figure S7), suggesting that some minor batch effects may be present despite stringent data quality filters and a common sequencing platform used for all samples. To minimise uncertainties arising from possible sequencing batch effects, our results focus on patterns of genetic structure that occur within batches, or which are genomically localised, and therefore robust to genome-wide differences in sequencing data.

Taxonomic analysis of raw sequencing reads with Kraken (Supp Figure S8) showed that with few exceptions all colonies harboured *Cladocopium* as the dominant symbiont. Further investigation of symbiont reads mapping to the *Cladocopium proliferum (syn. goreaui)*(*52*) genome revealed two distinct mitochondrial haplotypes (Figure 2; Supplementary Figure S9) and two major clusters based on the d2s kmer-based distance metric (Supplementary Figures S10). The geographic distribution of samples harbouring the two mitochondrial haplotypes suggests that they correspond to *Cladocopium* types, C1 and C2, previously identified using SSCP polymorphisms(*46*, *53*) in *A. kenti* at Magnetic Island (C1; 22/28 of MI samples) and Pelorus Island (C2; 18/30 of PI samples). While the C1 haplotype was strongly associated with three inshore reefs (MI, PR, DI) the C2 haplotype was more broadly distributed across inshore and offshore locations.

**Figure 2:**
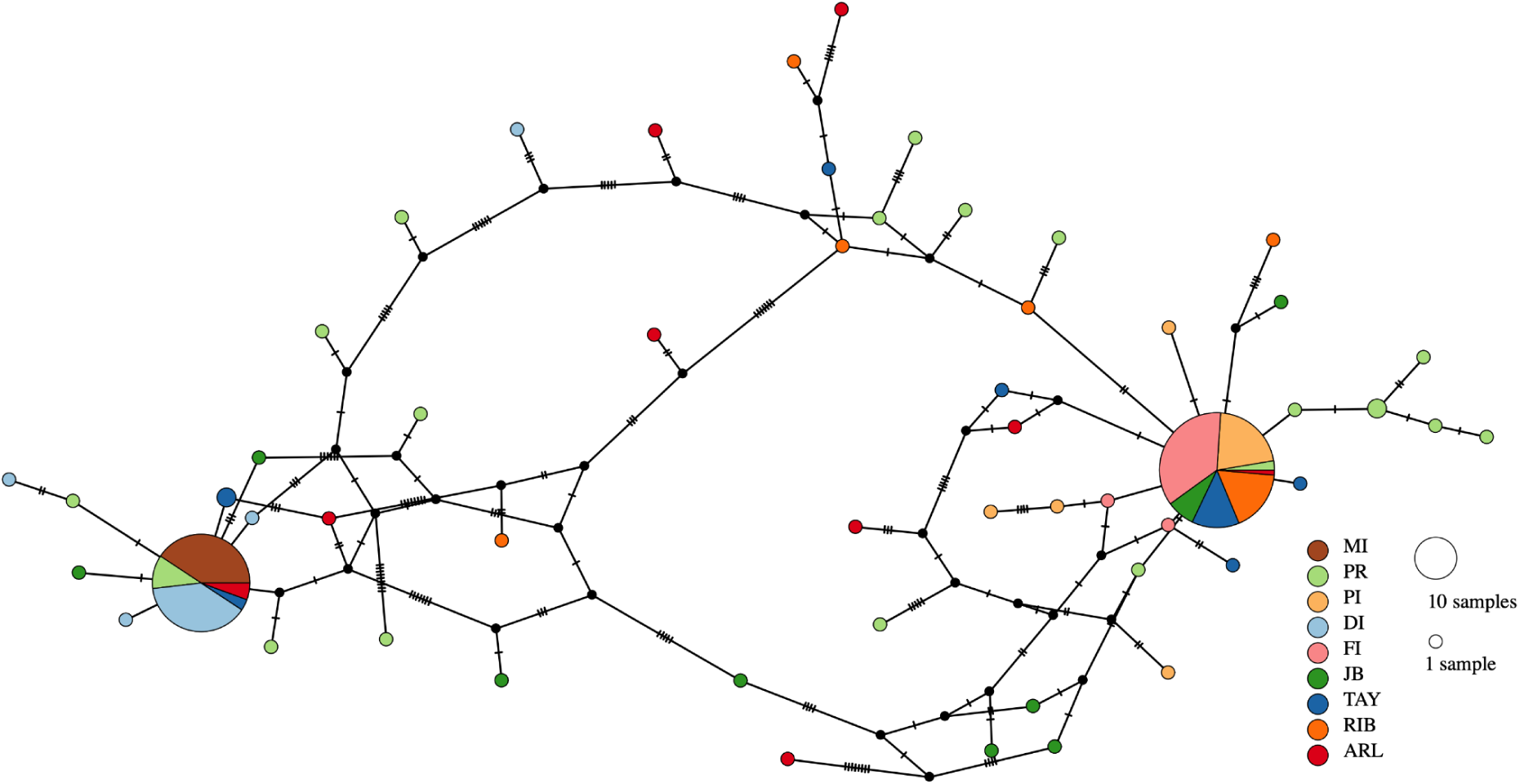
Symbiont mitochondrial haplotype network based on consensus sequences of reads mapping to the *Cladocopium proliferum (syn. goreaui)* mitochondrial genome within coral whole genome sequencing data. Individual haplotypes are shown as separate nodes with node size reflecting the number of samples. Edges connect related nodes with cross bars indicating the number of different sites. Sample location abbreviations and colours are the same as Figure 1.

### Identification of inversion loci and genotyping of individuals for inversion karyotypes

Absent or weak (see below) population structure among the eight non-Magnetic Island reefs allowed us to identify putative inversion polymorphisms using a scan for local genetic structure based on principal component analysis (Galinsky et al., 2016; Meisner et al., 2021). This scan (performed using PCAngsd on genotype likelihoods) revealed five genomic regions ranging in size from 200Kb to 2Mb exhibiting exceptionally strong population structure compared with the genomic background (Figure 3A). A similar scan for the Magnetic Island population failed to reveal any clear peaks in the strength of local population structure (Supplementary Figure S11), however, this probably reflects a lack of statistical power rather than absence of inversions at Magnetic Island (Supplementary Figure S11; Methods). All five genomic regions with strong local population structure in the non-Magnetic Island population also exhibited patterns of heterozygosity and linkage disequilibrium indicative of strongly suppressed recombination in heterokaryotypes, a feature often considered to be diagnostic for inversions(*23*, *24*), although chromosomal fusions and translocations may generate similar signatures(*54*). Firstly, visual inspection of population structure within each region revealed three major clusters along PC1 (Figure 3B; Supplementary Figure S12), as expected based on the three possible genotypes of an inversion polymorphism (Huang 2020, Harringmeyer 2022). Heterozygosity within each locus was highest in corals assigned to the central cluster (corresponding to heterokaryotypes) (Figure 3C; Supplementary Figure S12), which is an expected consequence of sharply reduced recombination between inverted alleles. Finally, linkage disequilibrium was elevated within the inverted region for heterokaryotypes (Figure 3D) but less so for homokaryotypes. This LD effect was clearly observable for putative inversions L1 and L2 but not for inversions L3-L5 likely due to their smaller sizes(0.2-0.5Mb) and occurrence in less-contiguous regions of the assembled genome (Supplementary Figure S13).

**Figure 3:**
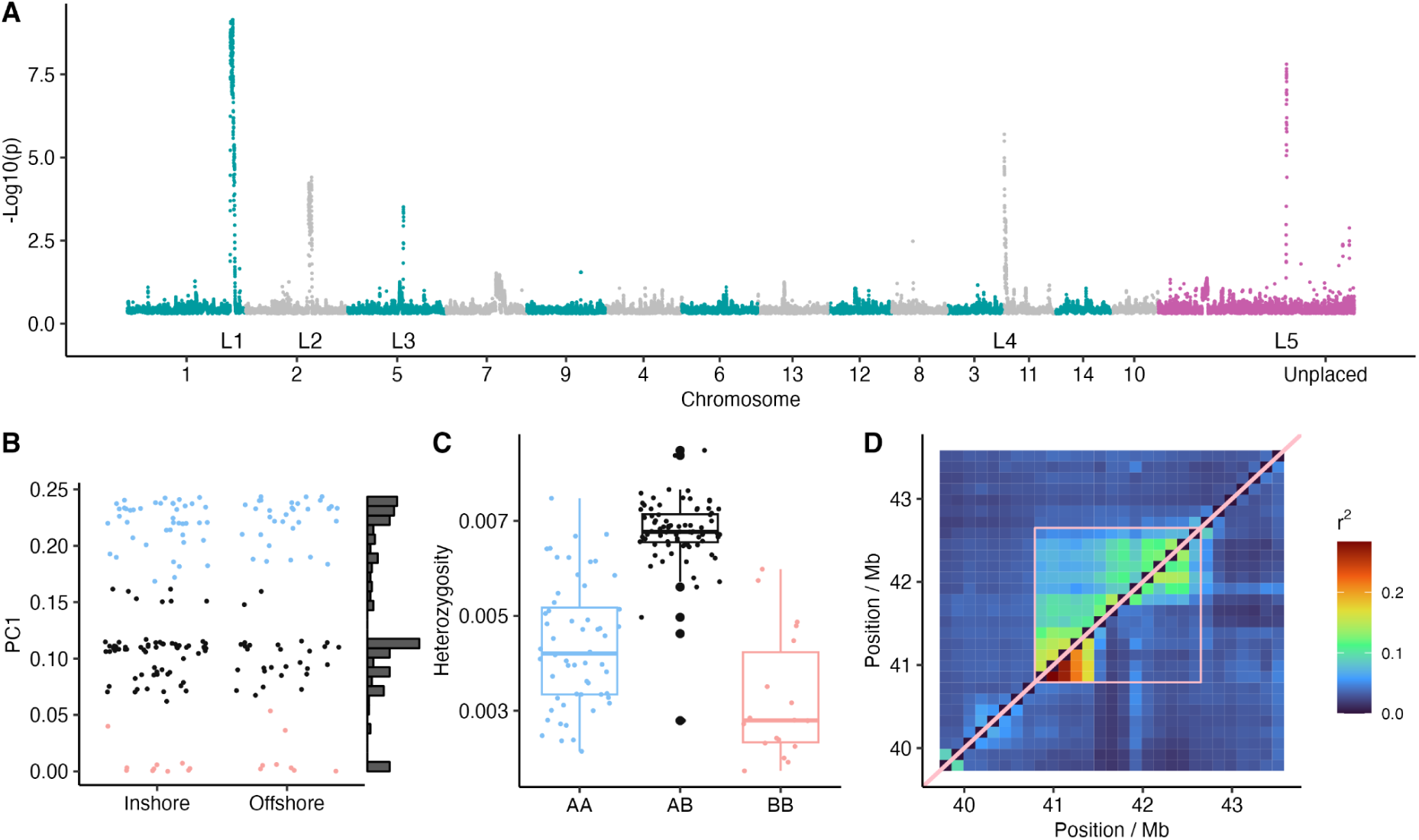
Location and characteristics of highly polymorphic inversions in *A. kenti*. **A:** Manhattan plot showing p-values derived from Galinsky statistics indicative of local population structure with five putative inversions (L1-L5) visible as strong signals compared with the background. Genomic coordinates reflect placement of *A. kenti* scaffolds via alignment to the *Acropora millepora* chromosome level genome. Alternating green and grey points indicate chromosome membership while pink points are on unplaced scaffolds. Plots **B-D** show the hallmarks of an inversion for L1 (similar plots for all other inversions are provided in supplementary figures S12 and S13). **B** shows strong local population structure with three clusters along PC1. **C** shows individual heterozygosity within the L1 region with individual corals genotyped for the L1 inversion according to their corresponding cluster in B. **D** shows pairwise linkage (r^2’^ statistic) across a 3Mb region centered on the L1 inversion. Top diagonal represents 93 heterozygous individuals and bottom diagonal 76 individuals homozygous for the major allele (AA). Each grid square shows the average value for all relevant SNPs. Pink box delineates inversion bounds inferred from PCAngsd (part A).

In addition to genetic evidence for inversions presented in Figure 3 we used the structural variant detection software Manta(*55*) to obtain direct read-based evidence for inversion events for two samples for which we had sequencing data at greater depth (∼20×). One of these deeper sequenced samples (FI-1-3) was from the non-Magnetic Island population, and the other from Magnetic Island (MI-1-4). PCA-based genotyping (eg Fig 3B) for the non-Magnetic Island individual indicated that it was heterozygous for the L1, L3 and L4 inversions, however, of these only the L1 inversion was contained within a single scaffold of our assembly and therefore suitable for analysis with Manta. In this individual, short-read data showed clear support for a 1.2Mb inversion event (called by Manta and manually verified; Supplementary figure S14) that closely matched the L1 region identified via local genetic structure (Figure 3). Although it was not possible to genotype the Magnetic Island individual (MI-1-4) using local PCA due to the small sample size at that location, short-read data confirmed that it was also heterozygous for the L1 inversion (Supplementary Figure S14).

All five inversion loci detectable via our local PCA-based method were present at high frequency (0.17-0.34) and were polymorphic across all reef sites (Figure 4A). None deviated significantly from genotype proportions expected under Hardy Weinberg equilibrium (p>0.19) as might be expected under strong spatially divergent selection (excess of homokaryotypes), or if recessive mutations resulted in a lethal homokaryotype (excess of heterokaryotypes). To investigate mutational load we used SnpEff(*56*) to predict the severity of impact of SNPs on protein coding genes. SnpEff predicts coding effects such as start/stop codon gains or losses, frameshifts and amino acid substitutions, and classifies these into categories ranging from low (e.g. synonymous variant) to high impact (e.g. frameshift, premature stop codon). Ignoring intergenic regions to avoid confounding effects from gene density, we compared the predicted impact of SNPs within inversions and contrasted this with an equal number of SNPs from a random selection of 100 × 50kb regions scattered throughout the genome. We found that the predicted impact of SNPs within inversions was shifted towards higher values compared with the genomic background (Figure 4B). Variants in the highest impact category are predicted to cause major disruptions to the protein sequence and are therefore likely to be associated with deleterious effects. These highly disruptive variants were more abundant in inversions across all allele frequency classes.

**Figure 4:**
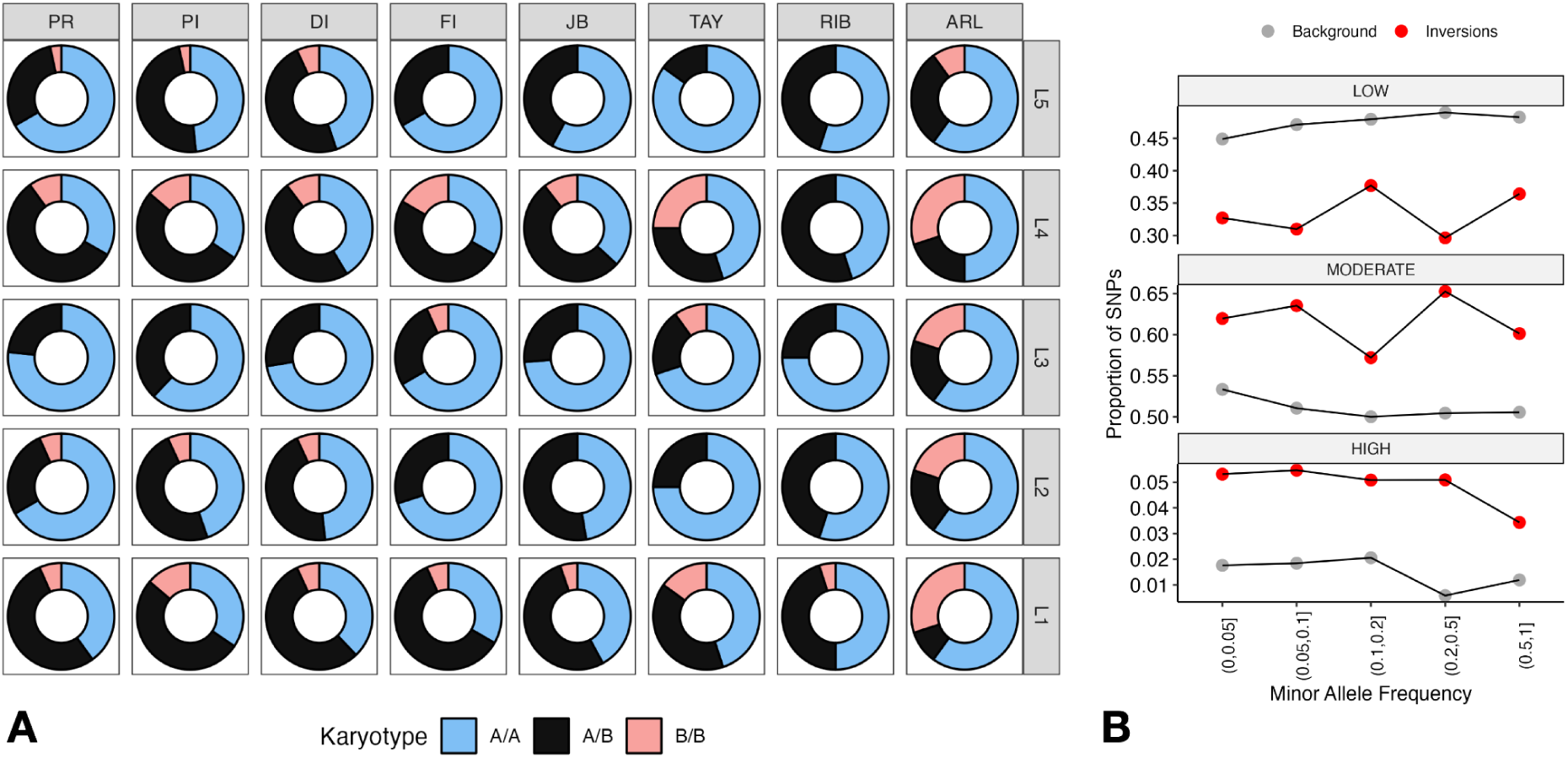
**A:** Distribution of karyotype frequencies for inversions across all non-Magnetic Island sites. The letter A is used to denote the most frequent karyotype. **B:** Mutational load of inversions in *A. kenti*. Proportions of SNPs rated as low, moderate and high impact by SnpEff are shown across all allele frequency classes within inversions (red) and in the background collinear genome (grey).

### Signatures of selection are not associated with inversion loci

Since inversions can facilitate divergent selection under gene flow(*19*, *27*) we identified two pairs of contrasting environmental conditions against which such divergence would be expected to occur in our study; (1) inshore versus offshore sites which differ strongly in nutrient, thermal and turbidity regimes(*48–50*), and (2) individuals clearly harbouring C1 or C2 dominant symbiont communities (most common haplotypes shown in Figure 2). Analyses with AMOVA failed to identify any significant associations between allele frequencies of any of the five inversions and reef, shore (inshore vs offshore) or dominant symbiont (Supplementary Table S5).

To identify sites potentially under divergent selection in the non-Magnetic Island population independent of inversions, we performed genome-wide scans of pairwise F_ST_ between inshore and offshore, and between C1 and C2 dominant colonies. We found that none of the five inversions overlapped with these highly divergent regions (F_ST_ z-score>6) and average F_ST_ within all inversions was within 3 standard deviations of the mean in all cases. In addition, levels of absolute divergence (D_xy_) were not elevated in these highly differentiated regions indicating that they are not regions of locally restricted gene flow (islands of speciation (*57*)) or inversions that we did not detect via our PCA-based method. Further analysis of highly differentiated regions in the non-Magnetic Island population revealed that they did not generally have the characteristics of strong selective sweeps (reduced Tajima’s D, reduced D_xy_; Figure 5B,5D).

**Figure 5:**
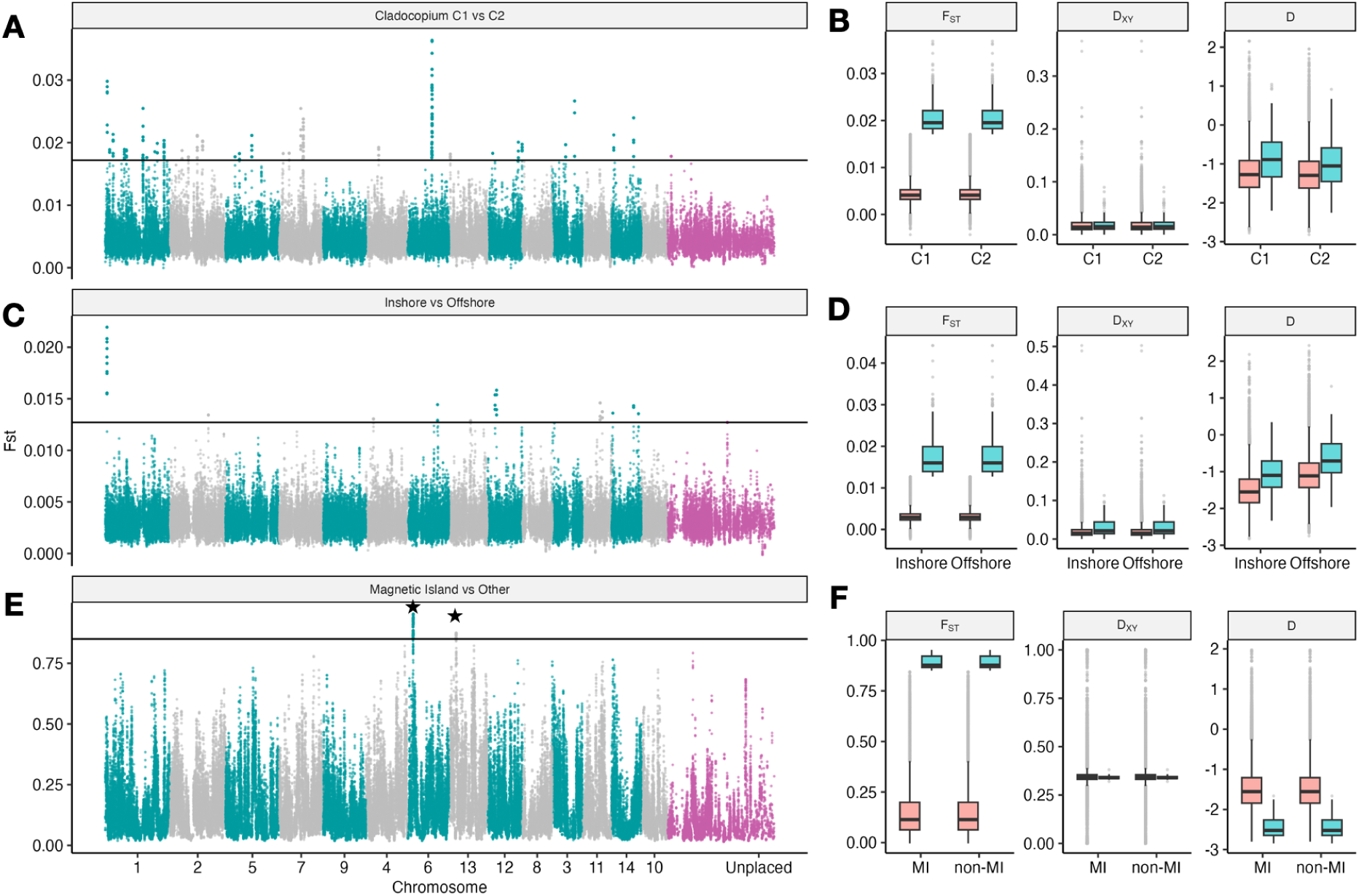
Genome-wide scan for highly differentiated regions between colonies dominated by C1 or C2 symbionts (A,B), inshore and offshore reefs (C,D) and Magnetic Island and non-Magnetic Island (E,F). Manhattan plots (A,C,E) show F_ST_ calculated within 20kb windows with the horizontal line delineating extreme (z-score > 6) values. Boxplots (B,D,F) to the right of each Manhattan plot show corresponding diversity statistics, F_ST_, D_xy_ and Tajima’s D calculated in extreme F_ST_ regions (blue) and contrasted with the genomic background (pink). Regions identified as putative selective sweeps are indicated with stars in E.

Despite a strong background level of divergence reflecting historical separation(*51*), we identified two exceptionally differentiated regions between Magnetic Island and non-Magnetic Island populations (Figure 5E). Tajima’s D in both these highly differentiated regions was sharply reduced (Figure 5F) compared with the genomic background indicating that divergence in these regions is associated with strong linked selection (selective sweeps). One of these regions overlaps with a selective sweep that was previously identified from inshore samples and contains a Tandem repeat of EGFR genes (*51*).

Inversions captured 214 genes with diverse functions, however, we found that there was a strong enrichment (p=8e-5; Fisher Exact Test in topGO) for genes involved in DNA-binding (GO:0003677) due to the presence of 14 genes with this GO term across the L1, L2 and L5 inversions. Of these, 4 were involved in transcriptional regulation and 3 were ATP-dependent helicases (1 on L1, 2 on L2), a protein family that plays crucial roles in recombination and DNA repair.

## Discussion

Our results clearly show that at least five chromosomal rearrangements, most likely inversions, exist as common polymorphisms within *Acropora kenti* on the central Great Barrier Reef. Elevated mutational load within these inversions and uniformly high frequencies of the minor karyotype among reefs (Figure 4) suggests they are under balancing selection. Inversions such as these (lacking strong spatial or ecological structure) are relatively challenging to detect because they do not produce signatures of elevated divergence (large blocks of high F_ST_) and can only be identified through direct read-based methods or local PCA (eg Figure 3). Moreover, datasets that include overall population structure (in the collinear genome) might fail to detect inversions through local PCA analysis as signals from inversions would be difficult to distinguish from background population structure. Our results also suggest that identifying inversions through local PCA analysis requires a large sample size as we were unable to detect any signatures of inversions at Magnetic Island (28 samples) despite direct read-based evidence from one individual (MI-1-4) that at least the L1 inversion is present in this population. These factors suggest that inversion polymorphisms of the type identified here may be present in many coral populations, but have been overlooked until now due to a historical focus on signatures of selection(*14*, *51*) and population structure(*8*, *58*) in coral genomic studies.

Our observation that mutational load (Figure 4B) was higher within inversions compared with the background genome is consistent with similar findings in butterflies(*33*) and sunflowers(*36*), as well as theoretical work(*35*) showing that reduced effective population size within inversions contributes to higher rates of genetic drift. An important consequence of accumulated mutational load within inversions is that if harmful mutations are completely or partially recessive the heterokaryotype will have higher fitness leading to a balanced polymorphism maintained by associative overdominance(*19*, *35*, *59*). Heterokaryotypes have higher fitness under associative overdominance because recessive mutations accumulate at different loci on each karyotype. In the case where these fitness differences are extremely strong (e.g. lethal before maturity) they may lead to detectable deviations from Hardy Weinberg equilibrium (HWE)(*33*, *34*). Therefore, since none of the five inversions identified in this paper deviated significantly from HWE (Figure 4A) they are unlikely to be associated with recessive lethal effects. However, given the large number of high-impact SNPs found within inversions by SnpEff some moderate fitness impacts are expected, and these may not have generated detectable deviations from HWE as doing so requires much larger sample sizes than used in our study (*60*).

Elevated mutational load at inversions in *A. kenti* may have important conservation genetics implications. In particular, the presence of recessive harmful loci in linkage increases the chances that population bottlenecks could result in inbreeding depression. This could be mitigated in conservation management programs such as coral aquaculture or assisted gene flow by screening colonies for spawning to ensure a diversity of inversion karyotypes. The potential role of inversions in local or clinal adaptation should also be considered. Although our study was unable to identify any ecological or spatial factors associated with adaptive selection at inversions, future studies at different spatial scales, or that measure different ecological variables should attempt to do so. This is important because inversions have often been shown to underpin local adaptation and genetic interventions such as assisted gene flow should seek to preserve locally adaptive variation.

Structural variants including inversions are often large and have a major impact on evolutionary processes(*54*) yet our ability to detect and study them remains limited. Our results highlight the fact that structural variants such as inversions are present in corals, and can have important impacts on genetic diversity and fitness. As technologies for structural variant detection improve it is therefore important that SVs are considered alongside SNPs as a key component of genetic variation.

## Methods

### Sample collection and sequencing

Offshore samples were collected in March 2017 from four locations in the central GBR under Great Barrier Reef Marine Park Authority collection permit G16/38488.1. A. Reefs for offshore samples were selected to approximately match the latitudes of four inshore locations for which sequencing data was available from a previous study (*51*). A fifth inshore location (Magnetic Island) was also included but since this is known to harbour a genetically diverged population no matching offshore reef was considered necessary. This resulted in a total of 228 samples, including 80 from offshore locations (Arlington Reef (ARL) n=20, Taylor Reef (TAY) n=20, Rib Reef (RIB) n=20, and John Brewer Reef (JB) n=20) and 148 from inshore reefs including Magnetic Island (Fitzroy Island (FI) n=30, Dunk Island (DI) n=30, Palm Islands (PI) n=30, and Pandora reef (PR) n=30, Magnetic Island (MI) n=28).

All samples were sequenced using the same sequencing protocol (100bp paired-end) and platform (Illumina HiSeq). Sequencing depth was generally shallow (2-5×) for most samples but two (FI-1-3, MI-1-4) had much deeper coverage (>20×). Sequencing coverage for offshore samples was slightly higher on average (4-5×) than inshore (2-3×). Mapping and coverage statistics for all samples are summarised in Supplementary Figure S1 and Supplementary Table S1.

### Data pre-processing and mapping

We followed the gatk germline variant calling best-practices workflow to generate mapped bam files from raw reads for each sample. Reads passing quality checks from each sample and lane were converted to unmapped bam format (uBAM) files. Adapters were marked using MarkIlluminaAdapters (Picard) before mapping to the reference genome assembly using bwa (v0.7.17-r1188). After mapping, PCR and optical duplicates were marked using MarkDuplicates (Picard). Two samples (FI-1-3, MI-1-4) sequenced at much higher coverage (28×, 26×), were downsampled using sambamba (v0.8.2) to achieve a target depth of approximately 3×.

### Removal of clones and misidentified samples

To ensure no cryptic species or misidentified samples were present in our data, we first reconstructed a mitochondrial genome for each sample by aligning raw reads to the reference mitogenome sequence of *Acropora kenti* (Genbank accession AF338425) and then extracting the most common base at each position using the -doFasta 2 option in ANGSD. These sequences of our samples were then used as queries to search the NCBI non-redundant nucleotide sequence database (nt June 23 2022) using megablast (v0.8.2) (Morgulis et al., 2008) with the option to output a maximum of five best matches with taxonomy information. While the best match for most samples was the mitogenome of *A. kenti*, nine samples from Arlington reef and one sample from John Brewer Reef matched *Acropora echinata* or *Acropora florida* (Supplementary Table S2). Next, we inferred a phylogenetic tree using IQ-TREE (v1.6.4) (Nguyen et al., 2015) based on the alignments (mafft v7.394) (Katoh & Standley, 2013) of mitogenome sequences of all samples together with the reference mitochondrial sequences of *A. kenti* (AF338425) and *A. echinata* (LC201841.1). The resulting tree (Supplementary Figure S2) revealed that nine samples from Arlington Reef and one from John Brewer Reef formed a distinct monophyletic clade together with the *A. echinata* mitogenome. Since these same samples also had particularly low mapping rates and genome coverage (Supplementary Figure S1), it is highly likely that they were misidentified in the field. We therefore excluded these samples from all further analyses.

Clones and closely related samples were identified by first estimating pairwise relatedness statistics with ngsRelate v2 (Korneliussen & Moltke, 2015) (https://github.com/ANGSD/NgsRelate). Pairs of samples with outlying relatedness were then identified using R1 vs R0, and R1 vs KING-robust kinship plots(*61*). This revealed eight pairs of closely related (expected kinship of 0.125) samples all of which were from Magnetic Island (Supplementary Figure S3). Retaining samples with higher depth where possible we then removed 6 samples to ensure that no close relatives were present in further analyses.

After removal of mis-identified samples and close relatives our final sample set contained 212 in total including 22 from MI, 30 each from inshore reefs PR, PI, FI and DI, 20 each from offshore reefs RIB, TAY, 19 from JB and 11 from ARL.

### Variant calling and genotype likelihood calculations

To account for the uncertainty of genotypes of each site due to low (2-5×) per-sample sequencing coverage, we used ANGSD to estimate genotype likelihoods. These genotype likelihoods formed the basis of all population genomic analyses and unless otherwise specified were generated as follows. ANGSD was run using the genotype likelihood (GL) model from GATK (-gl 2), inferring major and minor alleles from GL data (-doMajorMinor 1), estimating allele frequencies from GL data (-doMaf 1). Read data was filtered to include only bases with a quality score of at least 30 (-minQ 30) and reads with a mapping quality score of at least 30 (-minMapQ). SNPs were filtered to remove rare alleles (MAF >0.05), keeping only sites with p value<10^-6^ (-SNP_pval 1e-6) and only sites with data for at least 100 individuals.

Analysis with ANGSD was restricted to a 258Mb subset of the genome to avoid duplicated, low complexity and poorly assembled regions as follows. First, GENMAP (v1.2.0) (Pockrandt et al., 2020) was used to estimate the mappability of each site. Mappability scores were computed with *k-*mers with no more than 2 mismatches (-K 50 -E 2), and sites were retained if they had a mappability score equal to one which suggests they can be uniquely mapped. Second, we used mdust (v2006.10.17; default parameters) to identify and exclude low-complexity regions in the genome. Third, we excluded any sites from short (< 1Mb) scaffolds to reduce the influence of artefacts at the ends of fragmented reference sequences. Finally, since regions with very high or very low mapping depth are often associated with ambiguous mapping due to repeats, we removed sites with global depth across all samples with a depth higher or lower than the range’s 1% percentile (minimum 17×, maximum 1102×).

### Population structure and admixture

We used PCAngsd (v1.10)(*62*) to explore population structure and calculate admixture coefficients for all individuals. As input to PCAngsd we use genotype likelihoods calculated with ANGSD (see above) across the entire dataset (212 samples). The output covariance matrix estimated based on individual allele frequency was then used to compute eigenvalues and eigenvectors using the R package eigen and to generate PCA plots. PCAngsd was also used to automatically infer the best number of clusters (K=2, -admix_auto 10000) and perform admixture analysis. This revealed four highly admixed individuals (MI-1-1_S10, PI-1-16_S16, DI-2-4_S17, ARL_15_S69) that were excluded from further analyses. We also used NGAdmix to explore admixture with alternative numbers of clusters (K=2, K=3). The results for K=2 were qualitatively similar to PCAngsd and results for K=3 showed little support for a third cluster. Results for NGSAdmix with K=2 are shown in Figure 1.

To confirm that structure within the non-Magnetic Island population was not obscured by the presence of Magnetic Island samples we reran ANGSD and PCAngsd excluding all Magnetic Island samples and admixed samples. A PCA plot based on this analysis (Supplementary Figure S4) showed no visible structure between inshore and offshore.

To complement PCAngsd analyses we built sample trees based on pairwise genetic distances measured using the identity-by-state (-doIBS 1) calculation in ANGSD which randomly samples a single read from each position from each sample within filtered reference sites. The R function hclust was then used to generate a UPGMA tree from the IBS distance matrix and this was visualised with ggtree (*63*) (Supplementary Figure S5).

### Genome-wide estimates of genetic diversity within and between reefs

To calculate reef-specific diversity and divergence statistics, we first used the realSFS program within ANGSD to estimate one-dimensional (1D) folded site frequency spectra (SFS) for each of the nine reef locations separately, and two-dimensional folded SFS (2D) for each reef pair. Before estimating the SFS with realSFS, a two-step procedure was first implemented to generate a saf (site allele frequency likelihood) file followed by an optimisation of the saf file using ANGSD (-dosaf 1). Pairwise nucleotide diversity (π), Watterson’s θ, and Tajima’s D were estimated from the 1D-SFS of each reef using the thetaStat function within ANGSD with a sliding-window size of 10kb and s step size of 4kb. Global estimates of F_ST_ for each pair of reefs were computed directly from the 2D-SFS using the Reich estimator implemented in realSFS (Reich et al., 2009). A bootstrapped UPGMA tree based on pairwise F_ST_ was also generated using the R package ape (Supplementary Figure S6).

### Individual heterozygosity

The heterozygosity for each sample was estimated in ANGSD as the proportion of heterozygous sites in the 1D-SFS of each individual. A saf file was generated for each sample using ANGSD and used to estimate the 1D-SFS with the realSFS. The heterozygosity rate is calculated by dividing the number of variant sites by the total number of sites in R. This calculation was performed for each sample based on all available reads and then again for each sample after down-sampling to 2× coverage to determine whether differences in coverage could explain consistent differences between inshore and offshore samples (Supplementary Figure S7). The same procedure for calculating individual heterozygosity was also used within inversions by restricting the analysis to reads overlapping each inversion.

### Analysis of symbiont reads

To identify the dominant genus of Symbioiniaceae within each sample, we used the moqc pipeline (https://github.com/marine-omics/moqc) which performs a taxonomic classification of raw reads with the program KrakenUniq(*64*). The database for KrakenUniq included a representative full genome for each of the five major coral-associating genera as well as sequences from common contaminants and the genome of the coral host. Full details of all sequences used for database construction are provided as part of the moqc documentation.

Since KrakenUniq profiles for almost all samples showed *Cladocopium* as the dominant symbiont we mapped deduplicated reads to the *Cladocopium* mitochondrial genome (downloaded from http://symbs.reefgenomics.org/). We then used the doHaploCall function within ANGSD to obtain the consensus base at each position and print only positions where there is at least one variant allele (minMinor 1) and exclude positions where more than 10 individuals have an ambiguous base (maxMis 10). This resulted in alignment with 145 variable sites that we then cleaned further with goalign(*65*) to remove sequences from 29 samples that had more than 4% ambiguous bases. This alignment was used to generate a haplotype network with PopArt (Figure 2), and was also used to generate a maximum likelihood tree with IQ-Tree(*66*). IQ-Tree was run using ModelFinder to automatically detect the best evolutionary model and with 1000 ultrafast bootstraps. Visual inspection of this tree was used to identify individual samples belonging to the two most common haplogroups (Supplementary Figure S9) referred to in the text as C1 and C2.

To verify that distinctions between C1 and C2 symbiont harbouring colonies are not due to idiosyncrasies of mitochondrial genomes or difficulties calling consensus sequences from low coverage, we used d2ssect (https://github.com/bakeronit/d2ssect) to calculate pairwise distances between samples based on shared kmers within reads of symbiont origin using D2S statistic(*67*). All deduplicated reads that mapped to the Cladocopium genome were used for d2ssect analysis which generates a distance matrix based on d2s statistics. The cmdscale function in R was then used to reduce this distance matrix to 2 dimensions for plotting. The resulting plot, coloured by symbiont mito-haplogroup is shown in Supplementary Figure S10.

### Identifying and genotyping inversion loci

Inversion loci were initially identified using PCAngsd(*68*) as peaks in local genetic structure (Figure 3). PCAngsd was run on all non-hybrid non-Magnetic Island individuals (n=187) with the ‘-selection’ option which calculates variant weights based on the first principal component (Galinsky statistic). Results were converted into pseudo-chromosome coordinates using RagTag(*69*) (see below) and Galinsky statistics were converted to p-values using the pchisq function in R. Results for individual variants were then smoothed by calculating the average of -log10(p) within 100kb sliding windows with a 10kb step using bedtools(*70*). Inversion boundaries were then calculated by finding the start and end points of regions where -log10(p) exceeded 3 (p<0.001).

An attempt was made to replicate the inversion finding process described above for the 21 Magnetic Island individuals (excluding hybrids, clones and close-kin) but this did not yield any peaks exceeding our p-value threshold. To demonstrate that this lack of signal was most likely due to low sample size we performed the same analysis on a random subset of 21 individuals from the non-Magnetic Island population. The results of this analysis are shown in Supplementary Figure S11.

To infer inversion genotypes we first extracted genotype likelihood data specific to each inversion from the overall (genome-wide) genotype likelihoods calculated with ANGSD. A separate PCAngsd analysis was then run on each of these genotype likelihood files and genotypes were inferred by partitioning samples into three clusters along the first principal component (Supplementary Figure S12). The central cluster was always assumed to comprise heterozygotes (A/B) and clusters at the extremes were arbitrarily assigned to one of the two potential homozygous genotypes (A/A or B/B). Clusters were inferred using k-means clustering in R with k=3. Visual inspection of PC1 values and genotype assignments clearly indicates that not all samples could be unambiguously assigned to a single cluster, and it is therefore likely that our genotype assignments are not error-free. Nevertheless, we found that genotype proportions for all inversions did not deviate significantly from those expected under Hardy Weinberg Equilibrium (Supplementary Table S4)

### Heterozygosity, linkage disequilibrium and pairwise sample distances within inversions

To calculate individual heterozygosity within each inversion we used ANGSD and realSFS to generate a folded allele frequency spectrum for each sample and then calculated heterozygosity as the proportion of heterozygotes to invariant sites.

Linkage disequilibrium was calculated for genomic regions including each inversion plus up to 1Mb of flanking sequence at each end. Linkage disequilibrium statistics were then calculated using ngsLD(*71*) on genotype likelihood data for SNPs within each of these regions with individuals grouped into genotypes inferred via PCA (see section above). To reduce computation time and output file size a random sample of 1% of SNPs was used for inversions L1-L4 and a sample of 50% of SNPs used for L5. Only SNPs with data for at least 25 individuals were included. Outputs from ngsLD were then further processed in R to calculate average values of the EM r^2^ statistic for all SNPs in a regular 30×30 grid over each interval.

To investigate the age of inversions relative to the split between Magnetic Island and non-Magnetic Island populations we calculated genetic distances between pairs of individuals at inversion loci using ngsDist(*72*) https://github.com/fgvieira/ngsDist. Starting with ANGSD genotype likelihoods calculated on all 212 correctly identified unrelated individuals, we extracted data for SNPs contained within each inversion and also for a set of 100, 50kb regions randomly sampled from the genome. We then used ngsDist to calculate a pairwise genetic distance matrix for each of these data subsets, using the ‘--pairwise_del’ option to ensure that only sites with data for both individuals were used. We then used the ‘hclust’ package in R to generate a UPGMA tree based on pairwise distance matrices for each of these datasets. These trees are shown in Supplementary Figure S14.

### Variant severity in inversions

Since ANGSD variant calling focuses entirely on SNPs we used the bcftools mpileup variant caller to call SNPs and small indels for the purposes of variant severity analysis. As input to bcftools we used the same read alignment files that were used for ANGSD and we used the -q 20 and -Q 20 flags to ignore low quality (Phred<20) basecalls and alignments (MAPQ<20). We then filtered variant calls to retain only biallelic variants with at least a minor allele count of 2 and with quality scores greater than 30. Variants were also removed if they had more than 50% missing genotypes or a highly skewed allele balance (FS<20). The resulting vcf file was then split into a component that overlapped with inversions and a background component that overlapped with 100 randomly selected 50kb regions. We then ran snpEff(*56*) (version 5.2a) on each of these files to predict variant impacts, restricting the analysis to genic regions to avoid biases due possible differences in gene density between regions. Variants were then tabulated according to the top 3 severity levels (low, medium, high) and allele frequency. In general we found that variant count decreased as a function of allele frequency, however, a relatively large number of variants with frequencies close to 1 (>0.99) were present. We removed these variants as they may represent assembly or sequencing errors in the reference.

### Significance of ecological variables in structuring genetic variance in inversions

We used analysis of molecular variance (AMOVA)(*73*) to test the significance of various factors in structuring genetic variance at inversion loci. For this analysis we used inversion genotypes inferred using k-means clustering along PC1 (see above) and did not consider SNPs within inversion haplotypes. Analyses for each inversion were conducted separately with two models ∼shore/reef and ∼symbiont/reef where shore codes for inshore and offshore reef locations. Only colonies that could be clearly assigned to either C1 or C2 dominant symbionts were used for the symbiont analysis. AMOVA’s were calculated using the amova function in the poppr package and statistical significance of variance components were tested using the randtest.amova function with 999 random permutations. A summary of p-values and phi statistics for all tests is provided in Supplementary Table 2.5.

### Calculating sliding-window population genetic statistics

We used ANGSD to estimate genome-wide patterns of pairwise F_ST_, genetic diversity, and Tajima’s D. F_ST_ and D_xy_ were calculated for three pairwise contrasts between samples; 1) inshore and offshore reefs excluding Magnetic Island, 2) non-Magnetic Island and Magnetic Island and 3) samples that could be unambiguously assigned as harbouring C1 vs C2 symbionts.

For each sample grouping, realSFS was used to estimate the 1D SFS and then the 2D SFS for each pair of groups. Pairwise F_ST_ was then calculated in sliding windows using (realSFS fst stats2 -type 1) with a window size of 20kb and step of 4kb. For each sample grouping theta statistics were estimated using thetaStat within ANGSD (thetaStat do_stat -type 1) using the same sliding windows for F_ST_ scans. To avoid false signals resulting from window-based statistics dominated by very little data in the window, we excluded windows where the number of available sites (passing quality checks) was less than 10% of all sites.

We used a Perl script getDxy.pl (https://github.com/mfumagalli/ngsPopGen/blob/9ee3a6d5e733c1e248e81bfc21514b0527da967b/scripts/getDxy.pl) provided by the ngsPopGen toolset to calculate the D_XY_ for every site in the mafs files generated by ANGSD, non-bi-allelic sites were removed in the calculation. Per-site D_XY_ values were then grouped into sliding windows from F_ST_ estimates and the average value was assigned as the value for each window using Bedtools intersect and groupby.

### GO term enrichment of genes captured by inversions

To investigate genes captured by inversions we used bedtools to find all 214 genes that overlapped with inversion coordinates. Functional annotations for these genes, and all genes in the *A. kenti* genome were obtained from supplementary material in (*51*). Functional enrichment of genes within inversions was tested using topGO with the weight01 algorithm which reduces the likelihood of false positive enrichment in higher level terms. Statistical significance of enrichment was assessed using Fisher’s exact test and terms with p<0.01 are listed in Supplementary Table S6. For the most significantly enriched term (GO:0003677) supporting genes are listed in Supplementary Table S7

### Building a pseudo-chromosome reference

To facilitate the visualisation of genome-wide genetic statistics in manhattan plots, we used ragtag v2.0.1 (Alonge et al., 2019) to align the *Acropora kenti* genome to the *Acropora millepora* chromosome-level genome assembly (Fuller, Mocellin, et al., 2020) with default parameters. 488 out of 614 *A. tenuis* scaffolds were placed accordingly, comprising 94.3% of the assembly. The results were used to translate the base position in the original *A. kenti* assembly into the pseudo-chromosome level assembly for visualisation purposes.

### Read-based evidence for inversions

We ran manta(*55*) with default parameters to call structural variants from one deeply sequenced individual at Fitzroy Island and one at Magnetic Island. Inversions were extracted and overlapped using a consensus approach previously described (pubmed:31844586) and visualised using IGV (pubmed: 22517427) for manual validation.

Detailed methods including code and additional data is available on github https://github.com/bakeronit/acropora_kenti_wgs. Raw sequencing data is available under the NCBI Bioprojects XXXXX and XXXXX.

Supplementary Figures

Supplementary Tables

## References

1. L. B. DeFilippo, L. C. McManus, D. E. Schindler, M. L. Pinsky, M. A. Colton, H. E. Fox, E. W. Tekwa, S. R. Palumbi, T. E. Essington, M. M. Webster, Assessing the potential for demographic restoration and assisted evolution to build climate resilience in coral reefs. Ecol. Appl., e2650 (2022).

2. L. C. McManus, D. L. Forrest, E. W. Tekwa, D. E. Schindler, M. A. Colton, M. M. Webster, T. E. Essington, S. R. Palumbi, P. J. Mumby, M. L. Pinsky, Evolution and connectivity influence the persistence and recovery of coral reefs under climate change in the Caribbean, Southwest Pacific, and Coral Triangle. Glob. Chang. Biol. 27, 4307–4321 (2021).

3. M. V. Matz, E. A. Treml, B. C. Haller, Estimating the potential for coral adaptation to global warming across the Indo-West Pacific. Glob. Chang. Biol. 26, 3473–3481 (2020).

4. C. A. Logan, J. P. Dunne, C. M. Eakin, S. D. Donner, Incorporating adaptive responses into future projections of coral bleaching. Glob. Chang. Biol. 20, 125–139 (2014).

5. M. A. Colton, L. C. McManus, D. E. Schindler, P. J. Mumby, S. R. Palumbi, M. M. Webster, T. E. Essington, H. E. Fox, D. L. Forrest, S. R. Schill, F. J. Pollock, L. B. DeFilippo, E. W. Tekwa, T. E. Walsworth, M. L. Pinsky, Coral conservation in a warming world must harness evolutionary adaptation. Nat Ecol Evol (2022), doi:10.1038/s41559-022-01854-4.

6. K. Anthony, L. K. Bay, R. Costanza, J. Firn, J. Gunn, P. Harrison, A. Heyward, P. Lundgren, D. Mead, T. Moore, P. J. Mumby, M. J. H. van Oppen, J. Robertson, M. C. Runge, D. J. Suggett, B. Schaffelke, D. Wachenfeld, T. Walshe, New interventions are needed to save coral reefs. Nat Ecol Evol. 1, 1420–1422 (2017).

7. I. B. Baums, A. C. Baker, S. W. Davies, A. G. Grottoli, C. D. Kenkel, S. A. Kitchen, I. B. Kuffner, T. C. LaJeunesse, M. V. Matz, M. W. Miller, J. E. Parkinson, A. A. Shantz, Considerations for maximizing the adaptive potential of restored coral populations in the western Atlantic. Ecol. Appl. 29, e01978 (2019).

8. P. Bongaerts, I. R. Cooke, H. Ying, D. Wels, S. den Haan, A. Hernandez-Agreda, C. A. Brunner, S. Dove, N. Englebert, G. Eyal, S. Forêt, M. Grinblat, K. B. Hay, S. Harii, D. C. Hayward, Y. Lin, M. Mihaljević, A. Moya, P. Muir, F. Sinniger, P. Smallhorn-West, G. Torda, M. A. Ragan, M. J. H. van Oppen, O. Hoegh-Guldberg, Morphological stasis masks ecologically divergent coral species on tropical reefs. Curr. Biol. 31, 2286–2298.e8 (2021).

9. N. H. Rose, R. A. Bay, M. K. Morikawa, L. Thomas, E. A. Sheets, S. R. Palumbi, Genomic analysis of distinct bleaching tolerances among cryptic coral species. Proc. Biol. Sci. 288, 20210678 (2021).

10. J. T. Ladner, S. R. Palumbi, Extensive sympatry, cryptic diversity and introgression throughout the geographic distribution of two coral species complexes. Mol. Ecol. 21, 2224–2238 (2012).

11. V. Lukoschek, C. Riginos, M. J. H. van Oppen, Congruent patterns of connectivity can inform management for broadcast spawning corals on the Great Barrier Reef. Mol. Ecol. 25, 3065–3080 (2016).

12. A. M. A. Matias, I. Popovic, J. A. Thia, I. R. Cooke, G. Torda, V. Lukoschek, L. K. Bay, S. W. Kim, C. Riginos, Cryptic diversity and spatial genetic variation in the coral *Acropora tenuis* and its endosymbionts across the Great Barrier Reef. Evol. Appl. (2022), doi:10.1111/eva.13435.

13. J. P. Rippe, G. Dixon, Z. L. Fuller, Y. Liao, M. Matz, Environmental specialization and cryptic genetic divergence in two massive coral species from the Florida Keys Reef Tract. Mol. Ecol. 30, 3468–3484 (2021).

14. L. Thomas, J. N. Underwood, N. H. Rose, Z. L. Fuller, Z. T. Richards, L. Dugal, C. M. Grimaldi, I. R. Cooke, S. R. Palumbi, J. P. Gilmour, Spatially varying selection between habitats drives physiological shifts and local adaptation in a broadcast spawning coral on a remote atoll in Western Australia. Sci Adv. 8, eabl9185 (2022).

15. J. Zhang, Z. T. Richards, A. A. S. Adam, C. X. Chan, C. Shinzato, J. Gilmour, L. Thomas, J. M. Strugnell, D. J. Miller, I. Cooke, Evolutionary responses of a reef-building coral to climate change at the end of the last glacial maximum. Mol. Biol. Evol. (2022), doi:10.1093/molbev/msac201.

16. Z. L. Fuller, V. J. L. Mocellin, L. A. Morris, N. Cantin, J. Shepherd, L. Sarre, J. Peng, Y. Liao, J. Pickrell, P. Andolfatto, M. Matz, L. K. Bay, M. Przeworski, Population genetics of the coral Acropora millepora: Toward genomic prediction of bleaching. Science. 369 (2020), doi:10.1126/science.aba4674.

17. M. Kirkpatrick, How and why chromosome inversions evolve. PLoS Biol. 8 (2010), doi:10.1371/journal.pbio.1000501.

18. M. Wellenreuther, L. Bernatchez, Eco-Evolutionary Genomics of Chromosomal Inversions. Trends Ecol. Evol. 33, 427–440 (2018).

19. E. L. Berdan, N. H. Barton, R. Butlin, B. Charlesworth, R. Faria, I. Fragata, K. J. Gilbert, P. Jay, M. Kapun, K. E. Lotterhos, C. Mérot, E. Durmaz Mitchell, M. Pascual, C. L. Peichel, M. Rafajlović, A. M. Westram, S. W. Schaeffer, K. Johannesson, T. Flatt, How chromosomal inversions reorient the evolutionary process. J. Evol. Biol. (2023), doi:10.1111/jeb.14242.

20. M. Kapun, E. D. Mitchell, T. J. Kawecki, P. Schmidt, T. Flatt, An Ancestral Balanced Inversion Polymorphism Confers Global Adaptation. Mol. Biol. Evol. 40 (2023), doi:10.1093/molbev/msad118.

21. M. Joron, L. Frezal, R. T. Jones, N. L. Chamberlain, S. F. Lee, C. R. Haag, A. Whibley, M. Becuwe, S. W. Baxter, L. Ferguson, P. A. Wilkinson, C. Salazar, C. Davidson, R. Clark, M. A. Quail, H. Beasley, R. Glithero, C. Lloyd, S. Sims, M. C. Jones, J. Rogers, C. D. Jiggins, R. H. ffrench-Constant, Chromosomal rearrangements maintain a polymorphic supergene controlling butterfly mimicry. Nature. 477, 203–206 (2011).

22. J. Wang, Y. Wurm, M. Nipitwattanaphon, O. Riba-Grognuz, Y.-C. Huang, D. Shoemaker, L. Keller, A Y-like social chromosome causes alternative colony organization in fire ants. Nature. 493, 664–668 (2013).

23. C. Mérot, E. L. Berdan, H. Cayuela, H. Djambazian, A.-L. Ferchaud, M. Laporte, E. Normandeau, J. Ragoussis, M. Wellenreuther, L. Bernatchez, Locally Adaptive Inversions Modulate Genetic Variation at Different Geographic Scales in a Seaweed Fly. Mol. Biol. Evol. 38, 3953–3971 (2021).

24. O. S. Harringmeyer, H. E. Hoekstra, Chromosomal inversion polymorphisms shape the genomic landscape of deer mice. Nat Ecol Evol (2022), doi:10.1038/s41559-022-01890-0.

25. D. Schluter, L. H. Rieseberg, Three problems in the genetics of speciation by selection. Proc. Natl. Acad. Sci. U. S. A. 119, e2122153119 (2022).

26. A. Tigano, V. L. Friesen, Genomics of local adaptation with gene flow. Mol. Ecol. 25, 2144–2164 (2016).

27. M. Kirkpatrick, N. Barton, Chromosome inversions, local adaptation and speciation. Genetics. 173, 419–434 (2006).

28. K. Huang, R. L. Andrew, G. L. Owens, K. L. Ostevik, L. H. Rieseberg, Multiple chromosomal inversions contribute to adaptive divergence of a dune sunflower ecotype, doi:10.1101/829622.

29. L. Meyer, P. Barry, F. Riquet, A. Foote, C. D. Sarkissian, R. Cunha, C. Arbiol, F. Cerqueira, E. Desmarais, A. Bordes, N. Bierne, B. Guinand, P.-A. Gagnaire, “Divergence and gene flow history at two large chromosomal inversions involved in long-snouted seahorse ecotype formation.” bioRxiv (2023), p. 2023.07.04.547634.

30. R. Faria, K. Johannesson, R. K. Butlin, A. M. Westram, Evolving Inversions. Trends Ecol. Evol. 34, 239–248 (2019).

31. S. M. Schaal, B. C. Haller, K. E. Lotterhos, Inversion invasions: when the genetic basis of local adaptation is concentrated within inversions in the face of gene flow. Philos. Trans. R. Soc. Lond. B Biol. Sci. 377, 20210200 (2022).

32. T. Connallon, C. Olito, Natural selection and the distribution of chromosomal inversion lengths. Mol. Ecol. 31, 3627–3641 (2022).

33. P. Jay, M. Chouteau, A. Whibley, H. Bastide, H. Parrinello, V. Llaurens, M. Joron, Mutation load at a mimicry supergene sheds new light on the evolution of inversion polymorphisms. Nat. Genet. 53, 288–293 (2021).

34. D. Lindtke, K. Lucek, V. Soria-Carrasco, R. Villoutreix, T. E. Farkas, R. Riesch, S. R. Dennis, Z. Gompert, P. Nosil, Long-term balancing selection on chromosomal variants associated with crypsis in a stick insect. Mol. Ecol. 26, 6189–6205 (2017).

35. E. L. Berdan, A. Blanckaert, R. K. Butlin, C. Bank, Deleterious mutation accumulation and the long-term fate of chromosomal inversions. PLoS Genet. 17, e1009411 (2021).

36. K. Huang, K. L. Ostevik, C. Elphinstone, M. Todesco, N. Bercovich, G. L. Owens, L. H. Rieseberg, Mutation Load in Sunflower Inversions Is Negatively Correlated with Inversion Heterozygosity. Mol. Biol. Evol. 39 (2022), doi:10.1093/molbev/msac101.

37. D. K. Fabian, M. Kapun, V. Nolte, R. Kofler, P. S. Schmidt, C. Schlötterer, T. Flatt, Genome-wide patterns of latitudinal differentiation among populations of Drosophila melanogaster from North America. Mol. Ecol. 21, 4748–4769 (2012).

38. A. R. Anderson, A. A. Hoffmann, S. W. McKechnie, P. A. Umina, A. R. Weeks, The latitudinal cline in the In(3R)Payne inversion polymorphism has shifted in the last 20 years in Australian Drosophila melanogaster populations. Mol. Ecol. 14, 851–858 (2005).

39. D. Ayala, R. F. Guerrero, M. Kirkpatrick, Reproductive isolation and local adaptation quantified for a chromosome inversion in a malaria mosquito. Evolution. 67, 946–958 (2013).

40. A. P. Fuentes-Pardo, E. D. Farrell, M. E. Pettersson, C. G. Sprehn, L. Andersson, The genomic basis and environmental correlates of local adaptation in the Atlantic horse mackerel (Trachurus trachurus). Evol. Appl. 16, 1201–1219 (2023).

41. M. Todesco, G. L. Owens, N. Bercovich, J.-S. Légaré, S. Soudi, D. O. Burge, K. Huang, K. L. Ostevik, E. B. M. Drummond, I. Imerovski, K. Lande, M. A. Pascual-Robles, M. Nanavati, M. Jahani, W. Cheung, S. E. Staton, S. Muños, R. Nielsen, L. A. Donovan, J. M. Burke, S. Yeaman, L. H. Rieseberg, Massive haplotypes underlie ecotypic differentiation in sunflowers. Nature. 584, 602–607 (2020).

42. N. S. Locatelli, S. A. Kitchen, K. H. Stankiewicz, C. Cornelia Osborne, Z. Dellaert, H. Elder, B. Kamel, H. R. Koch, N. D. Fogarty, I. B. Baums, 1 Genome assemblies and genetic maps highlight chromosome-scale macrosynteny in Atlantic 2 acroporids, doi:10.1101/2023.12.22.573044.

43. A. A. Hoffmann, L. H. Rieseberg, Revisiting the Impact of Inversions in Evolution: From Population Genetic Markers to Drivers of Adaptive Shifts and Speciation? Annu. Rev. Ecol. Evol. Syst. 39, 21–42 (2008).

44. T. C. L. Bridge, P. F. Cowman, A. M. Quattrini, V. E. Bonito, F. Sinniger, S. Harii, C. E. I. Head, J. Y. Hung, T. Halafihi, T. Rongo, A. H. Baird, A tenuis relationship: traditional taxonomy obscures systematics and biogeography of the “Acropora tenuis” (Scleractinia: Acroporidae) species complex. Zool. J. Linn. Soc. (2023), doi:10.1093/zoolinnean/zlad062.

45. K. L. E. Berry, M. O. Hoogenboom, F. Flores, A. P. Negri, Simulated coal spill causes mortality and growth inhibition in tropical marine organisms. Sci. Rep. 6, 25894 (2016).

46. D. Abrego, M. J. H. VAN Oppen, B. L. Willis, Onset of algal endosymbiont specificity varies among closely related species of Acropora corals during early ontogeny. Mol. Ecol. 18, 3532–3543 (2009).

47. K. M. Quigley, M. J. H. van Oppen, Predictive models for the selection of thermally tolerant corals based on offspring survival. Nat. Commun. 13, 1543 (2022).

48. M. J. Furnas, Catchments and Corals: Terrestrial Runoff to the Great Barrier Reef (Australian Institute of Marine Science, 2003).

49. S. L. Coles, P. L. Jokiel, “Effects of salinity on coral reefs” in Pollution in Tropical Aquatic Systems, D. W. Connell, D. W. Hawker, Eds. (CRC Press, Florida, 1992), pp. 147–166.

50. J. E. Brodie, M. Devlin, D. Haynes, J. Waterhouse, Assessment of the eutrophication status of the Great Barrier Reef lagoon (Australia). Biogeochemistry. 106, 281–302 (2011).

51. I. Cooke, H. Ying, S. Forêt, P. Bongaerts, J. M. Strugnell, O. Simakov, J. Zhang, M. A. Field, M. Rodriguez-Lanetty, S. C. Bell, D. G. Bourne, M. J. van Oppen, M. A. Ragan, D. J. Miller, Genomic signatures in the coral holobiont reveal host adaptations driven by Holocene climate change and reef specific symbionts. Sci Adv. 6 (2020), doi:10.1126/sciadv.abc6318.

52. C. C. Butler, K. E. Turnham, A. M. Lewis, M. R. Nitschke, M. E. Warner, D. W. Kemp, O. Hoegh-Guldberg, W. K. Fitt, M. J. H. van Oppen, T. C. LaJeunesse, Formal recognition of host-generalist species of dinoflagellate (Cladocopium, Symbiodiniaceae) mutualistic with Indo-Pacific reef corals. J. Phycol. 59, 698–711 (2023).

53. K. M. Quigley, C. Alvarez Roa, G. Torda, D. G. Bourne, B. L. Willis, Co-dynamics of Symbiodiniaceae and bacterial populations during the first year of symbiosis with Acropora tenuis juveniles. Microbiologyopen. 9, e959 (2020).

54. C. Mérot, R. A. Oomen, A. Tigano, M. Wellenreuther, A Roadmap for Understanding the Evolutionary Significance of Structural Genomic Variation. Trends Ecol. Evol. 35, 561–572 (2020).

55. X. Chen, O. Schulz-Trieglaff, R. Shaw, B. Barnes, F. Schlesinger, M. Källberg, A. J. Cox, S. Kruglyak, C. T. Saunders, Manta: rapid detection of structural variants and indels for germline and cancer sequencing applications. Bioinformatics. 32, 1220–1222 (2016).

56. P. Cingolani, A. Platts, L. L. Wang, M. Coon, T. Nguyen, L. Wang, S. J. Land, X. Lu, D. M. Ruden, A program for annotating and predicting the effects of single nucleotide polymorphisms, SnpEff: SNPs in the genome of Drosophila melanogaster strain w1118; iso-2; iso-3. Fly . 6, 80–92 (2012).

57. T. E. Cruickshank, M. W. Hahn, Reanalysis suggests that genomic islands of speciation are due to reduced diversity, not reduced gene flow. Mol. Ecol. 23, 3133–3157 (2014).

58. C. Shinzato, S. Mungpakdee, N. Arakaki, N. Satoh, Genome-wide SNP analysis explains coral diversity and recovery in the Ryukyu Archipelago. Sci. Rep. 5, 18211 (2015).

59. P. Pamilo, S. Pálsson, Associative overdominance, heterozygosity and fitness. Heredity. 81 (Pt 4), 381–389 (1998).

60. J. Lachance, Detecting selection-induced departures from Hardy-Weinberg proportions. Genet. Sel. Evol. 41, 15 (2009).

61. R. K. Waples, A. Albrechtsen, I. Moltke, Allele frequency-free inference of close familial relationships from genotypes or low-depth sequencing data. Mol. Ecol. 28, 35–48 (2019).

62. J. Meisner, A. Albrechtsen, K. Hanghøj, Detecting Selection in Low-Coverage High-Throughput Sequencing Data using Principal Component Analysis. bioRxiv (2021), p. 2021.03.01.432540.

63. G. Yu, D. K. Smith, H. Zhu, Y. Guan, T. T.-Y. Lam, Ggtree : An r package for visualization and annotation of phylogenetic trees with their covariates and other associated data. Methods Ecol. Evol. 8, 28–36 (2017).

64. F. P. Breitwieser, D. N. Baker, S. L. Salzberg, KrakenUniq: confident and fast metagenomics classification using unique k-mer counts. Genome Biol. 19, 198 (2018).

65. F. Lemoine, O. Gascuel, Gotree/Goalign: toolkit and Go API to facilitate the development of phylogenetic workflows. NAR Genom Bioinform. 3, lqab075 (2021).

66. L.-T. Nguyen, H. A. Schmidt, A. von Haeseler, B. Q. Minh, IQ-TREE: a fast and effective stochastic algorithm for estimating maximum-likelihood phylogenies. Mol. Biol. Evol. 32, 268–274 (2015).

67. K. Song, J. Ren, G. Reinert, M. Deng, M. S. Waterman, F. Sun, New developments of alignment-free sequence comparison: measures, statistics and next-generation sequencing. Brief. Bioinform. 15, 343–353 (2014).

68. J. Meisner, A. Albrechtsen, K. Hanghøj, Detecting selection in low-coverage high-throughput sequencing data using principal component analysis. BMC Bioinformatics. 22, 470 (2021).

69. M. Alonge, S. Soyk, S. Ramakrishnan, X. Wang, S. Goodwin, F. J. Sedlazeck, Z. B. Lippman, M. C. Schatz, RaGOO: fast and accurate reference-guided scaffolding of draft genomes. Genome Biol. 20, 224 (2019).

70. A. R. Quinlan, I. M. Hall, BEDTools: a flexible suite of utilities for comparing genomic features. Bioinformatics. 26, 841–842 (2010).

71. E. A. Fox, A. E. Wright, M. Fumagalli, F. G. Vieira, ngsLD: evaluating linkage disequilibrium using genotype likelihoods. Bioinformatics. 35, 3855–3856 (2019).

72. F. G. Vieira, F. Lassalle, T. S. Korneliussen, M. Fumagalli, Improving the estimation of genetic distances from Next-Generation Sequencing data. Biol. J. Linn. Soc. Lond. 117, 139–149 (2016).

73. L. Excoffier, P. E. Smouse, J. M. Quattro, Analysis of molecular variance inferred from metric distances among DNA haplotypes: application to human mitochondrial DNA restriction data. Genetics. 131, 479–491 (1992).

