## Supplementary Figures for "Chromosomal inversions harbour excess mutational load in the coral, *Acropora kenti,* on the Great Barrier Reef"

### Supplementary Materials

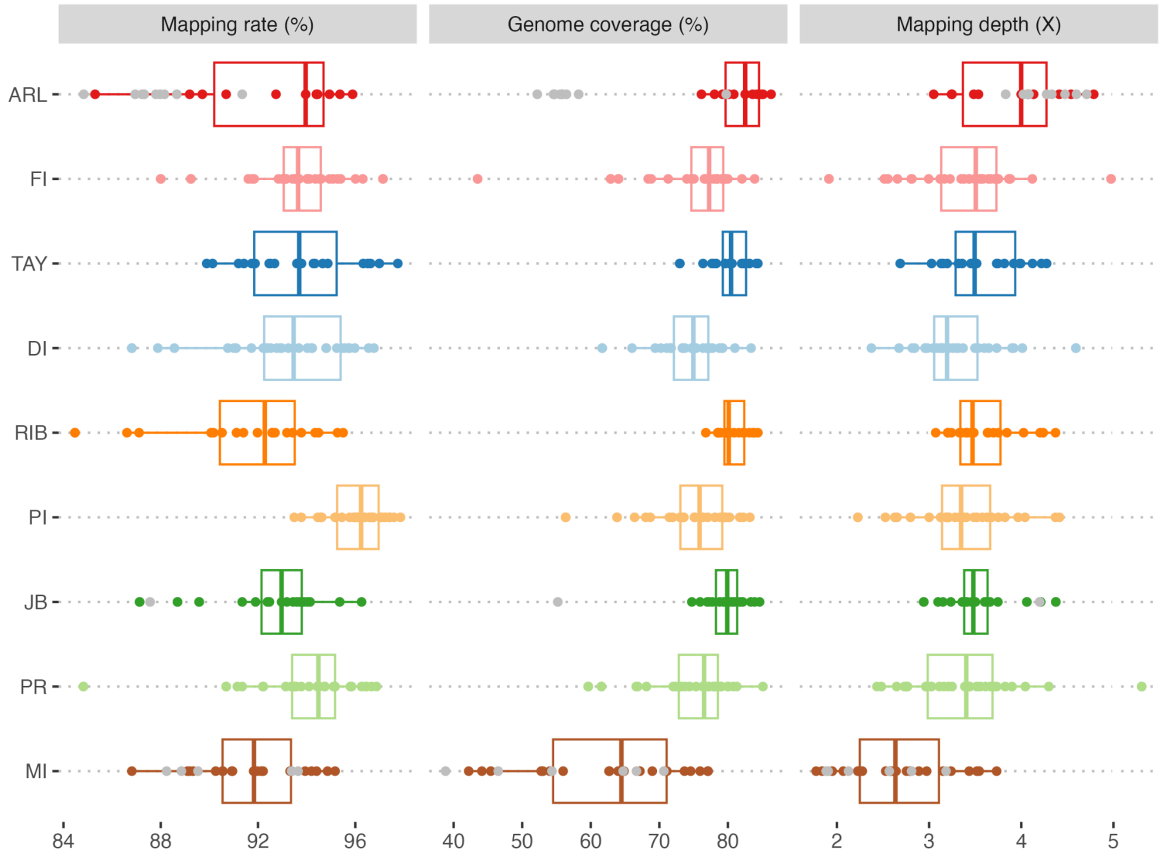


**Supplementary Figure S1**: Summary of read mapping statistics for all 228 *A. kenti* samples across 9 reef locations. Mapping rate refers to the proportion of all reads that could be successfully mapped to the *A. kenti* genome. Genome coverage is the proportion of the *A. kenti* genome covered by at least one read (mappable regions) and Mapping depth is the average depth of coverage within mappable regions.  Samples excluded from further analysis due to being species mis-ids or close kin are shown in grey.

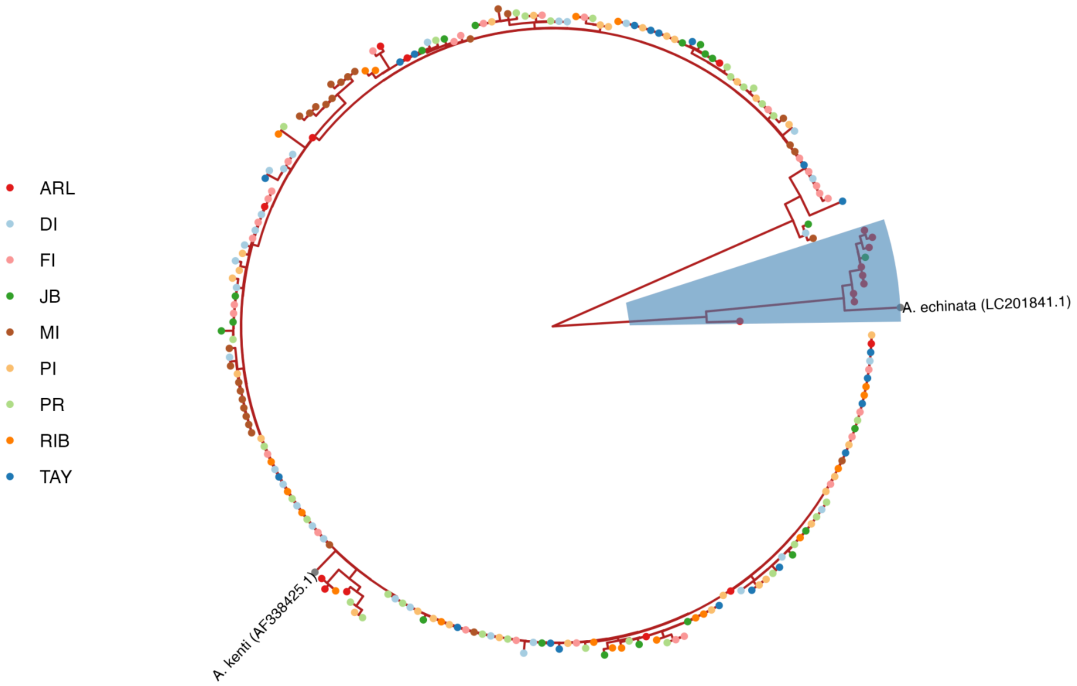


**Supplementary Figure S2**: Maximum likelihood tree of consensus mitochondrial genome sequences for all samples. Tip labels are coloured by reef location with the exception of reference sequences for *A. echinata* (LC201841.1) and *A. kenti* (AF338425.1) which are shown in grey. Samples within the highlighted clade (10 samples) were excluded from further analyses.

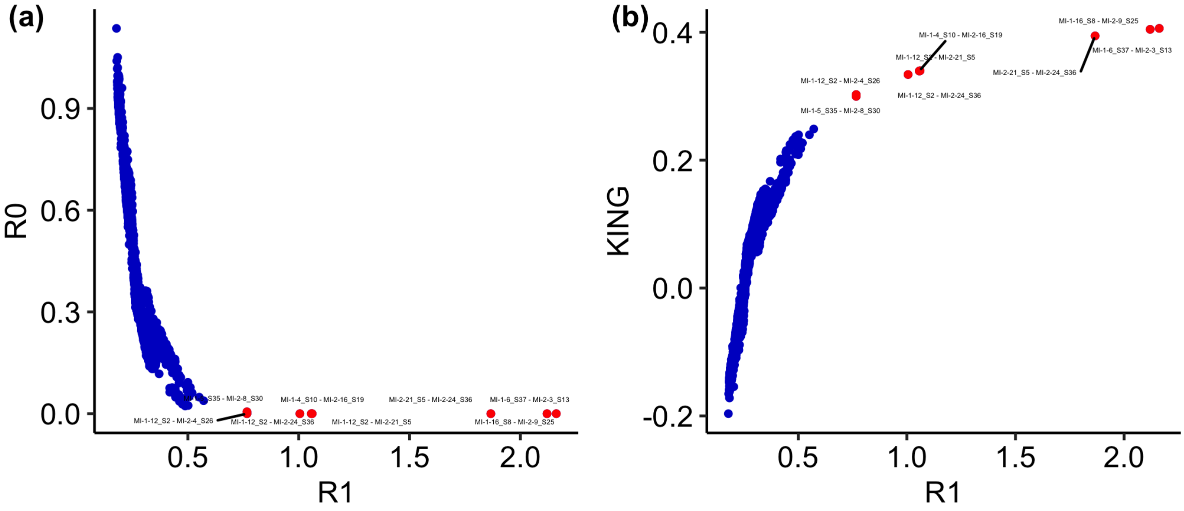

**Supplementary Figure S3**: Pairwise relatedness metrics for all pairs of samples: R0, R1 and KING statistics calculated with ngsRelate for all 218 *A. tenuis* samples. Pairs shown in red represent likely close-kin relationships and were used to select 6 samples for removal.

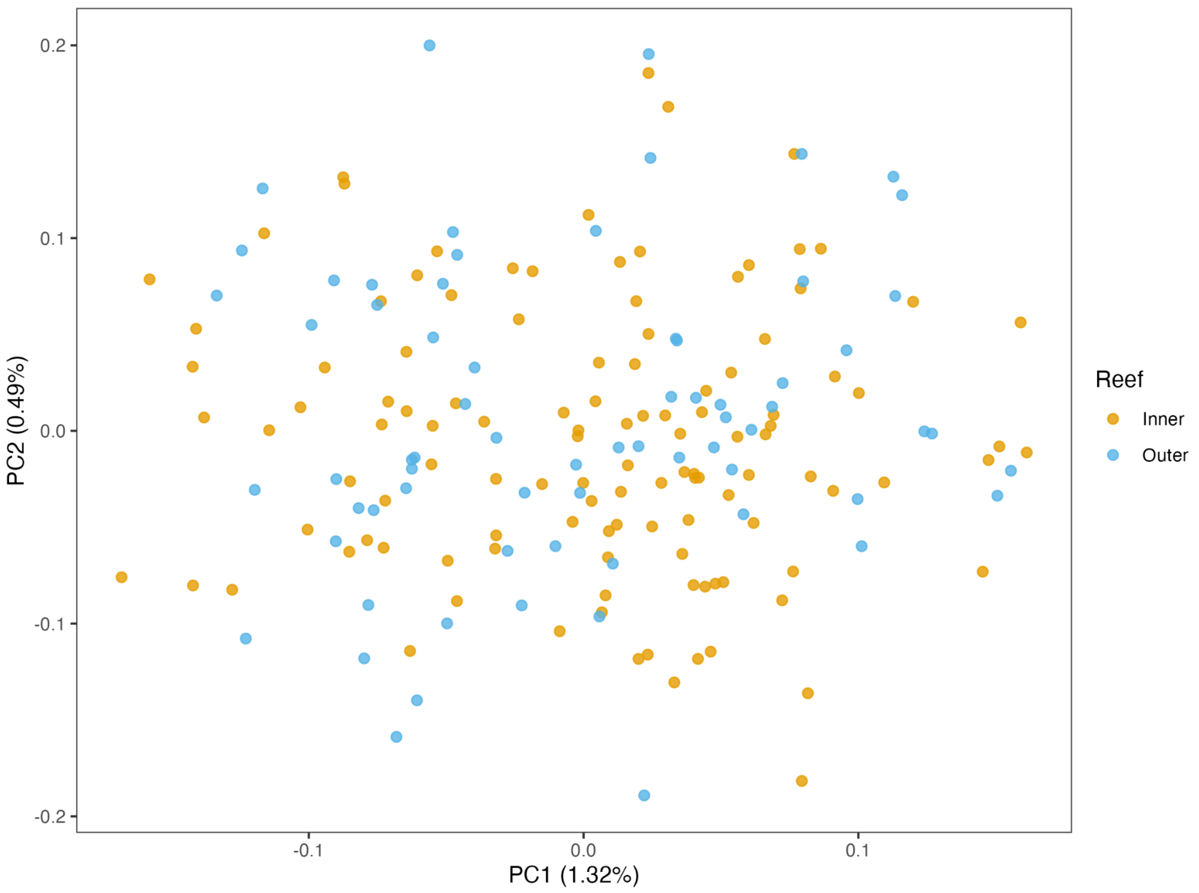


**Supplementary Figure S4:** Principal components analysis based on covariance matrix calculated with PCAngsd on non-Magnetic Island samples only. Samples labelled “inner” are from inshore reefs (PI, FI, PR, DI) and were sequenced in the first batch, while those labelled “outer” are from offshore reefs (ARL, TAY, RIB, JB) and were sequenced in the second batch.

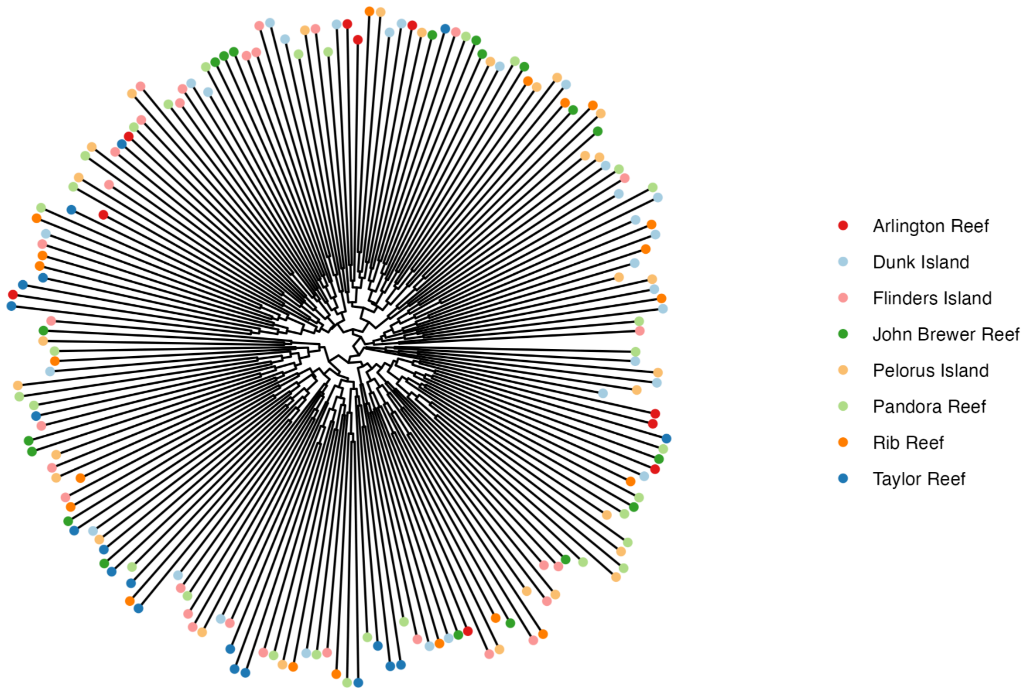


**Supplementary Figure S5:** Dendrogram built from pairwise identity by state (IBS) distances between all non-Magnetic Island samples.

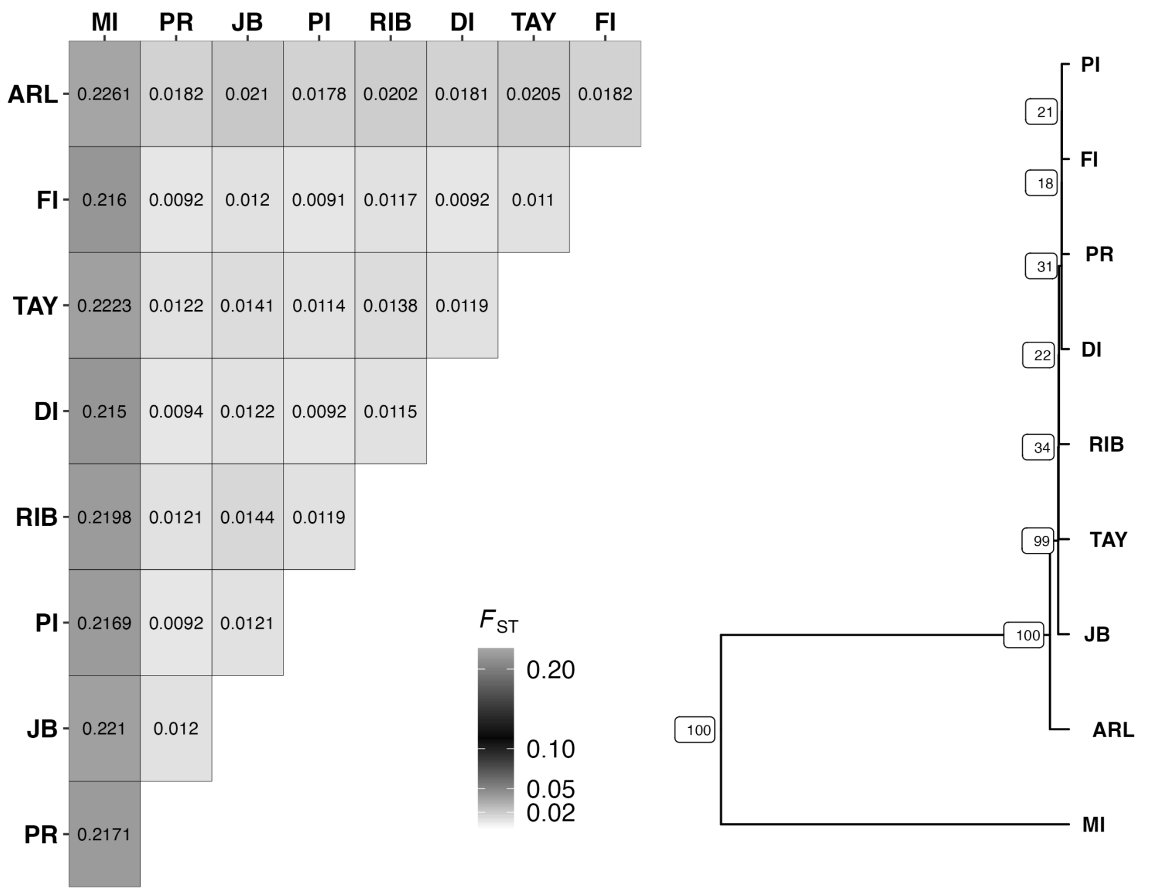


**Supplementary Figure S6:** Visualisation of pairwise Fst values between all nine sampling locations. Tree shown on the right is based on hierarchical clustering of the Fst distance matrix. Bootstrap support values are shown as node labels.

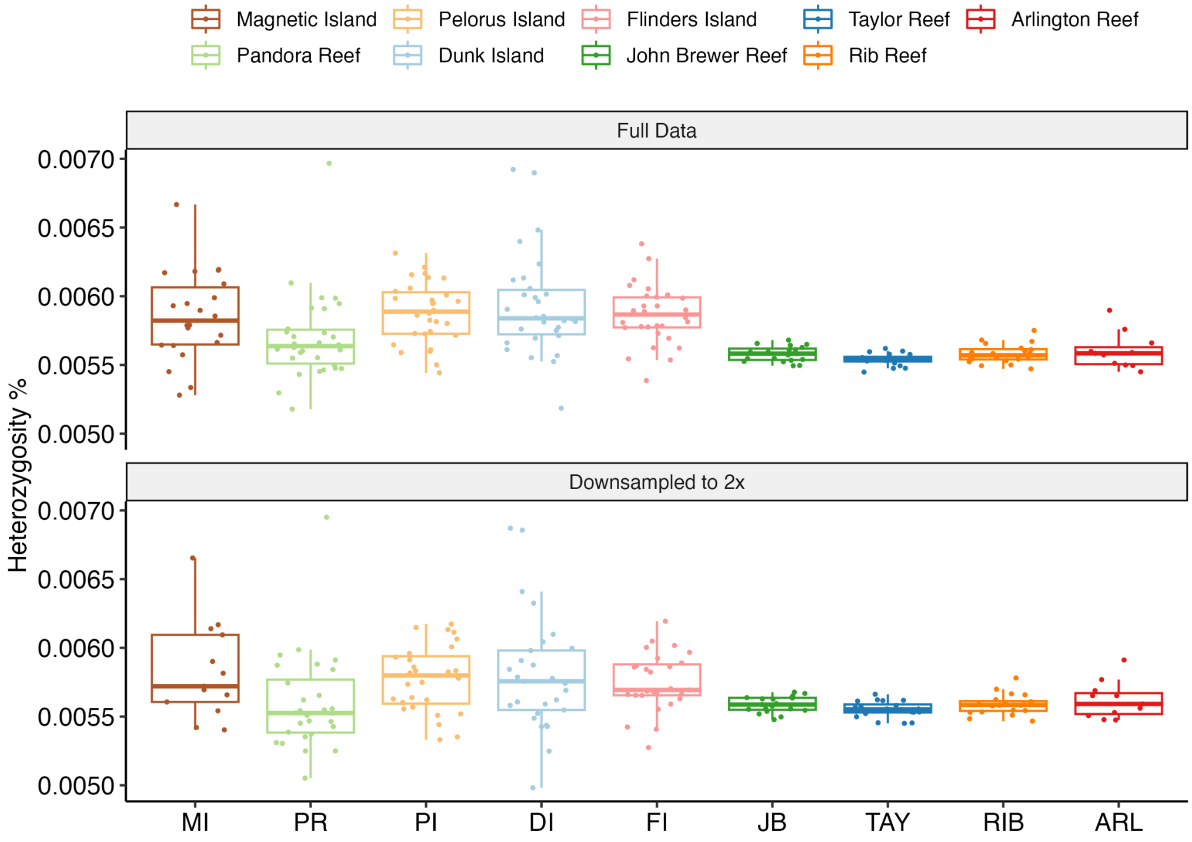


**Supplementary Figure S7**: Individual heterozygosity of samples at each reef location. Boxplots and individual points are shown for each reef for the full dataset (no downsampling) and for data that has been downsampled to 2x to achieve even coverage across all samples.

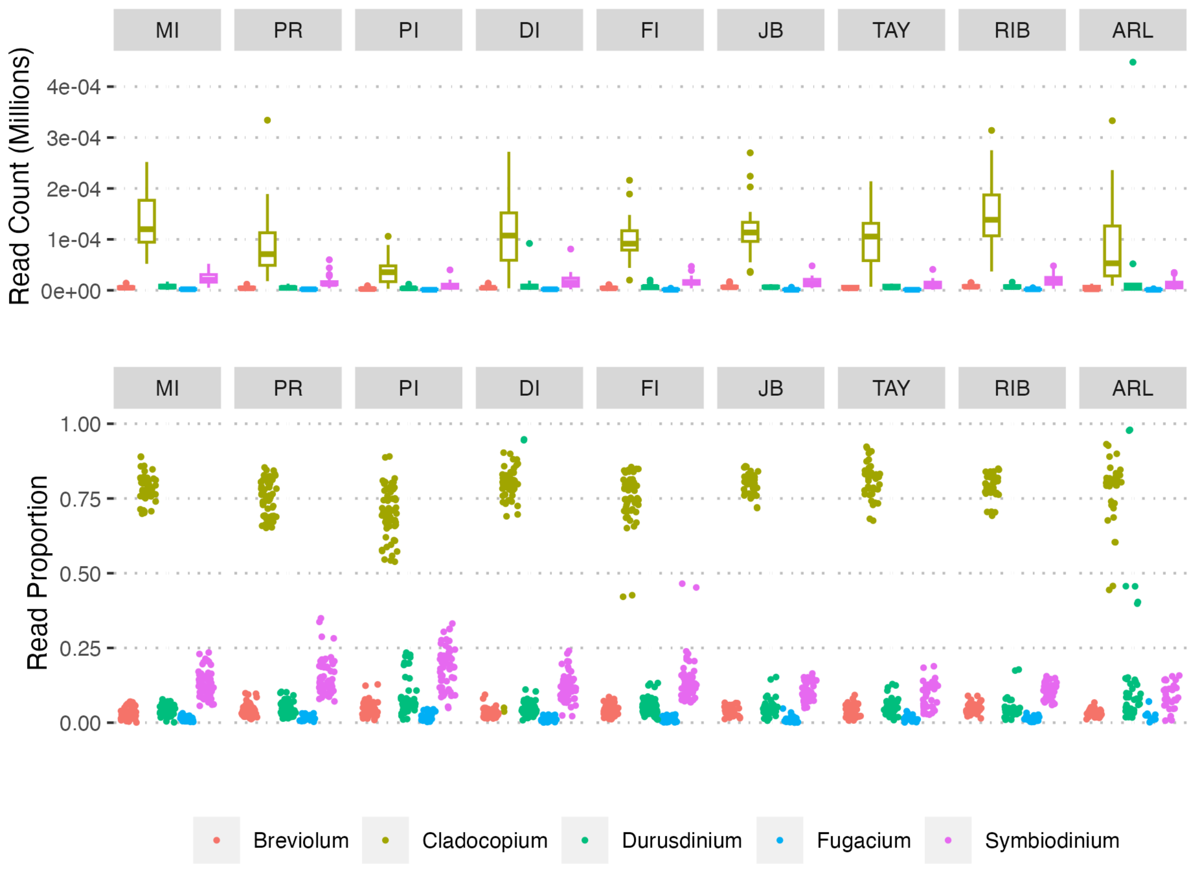


**Supplementary Figure S8:** Genus level profiles of Symbiodiniaceae present in whole genome sequencing data of *A. kenti* . Read counts represent numbers of reads classified by kraken as belonging to the given genus. Read proportions are relative to reads assigned to the family Symbiodiniaceae only.

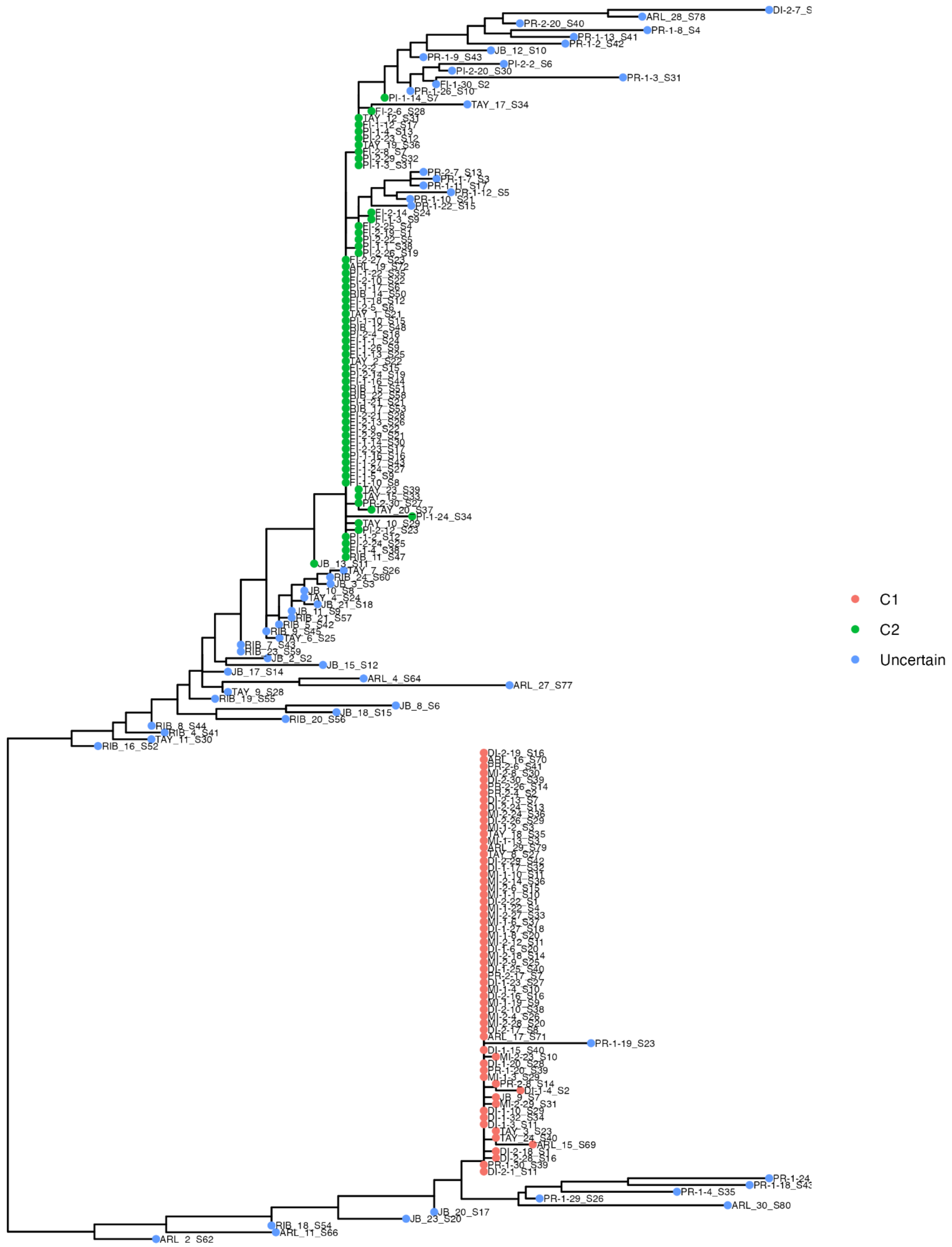


**Supplementary Figure S9:** Maximum likelihood tree of consensus symbiont mitochondrial genome sequences obtained from 189 samples for which sufficient sequencing data was available. Designation as C1 or C2 was made manually for samples with identical or near identical sequences belonging to two large haplogroups. Assignment of these haplogroups as C1 and C2 is based on locations of samples following results of Abrego et al 2009(Abrego et al. 2009)

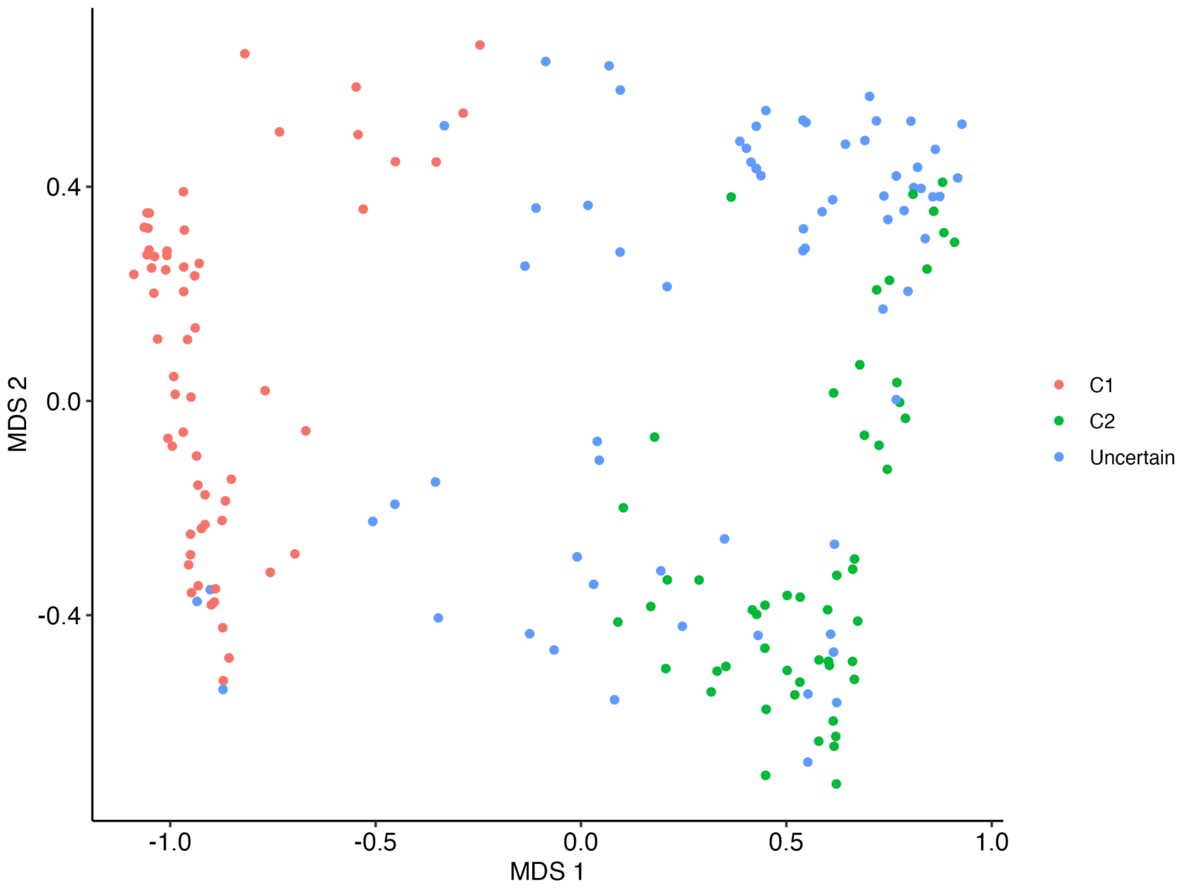


**Supplementary Figure S10:** Multidimensional Scaling (MDS) plot based on d2s distances calculated from kmer profiles of all reads that map to the Cladocopium genome. Designation as C1 or C2 is the same as for Figure S9.

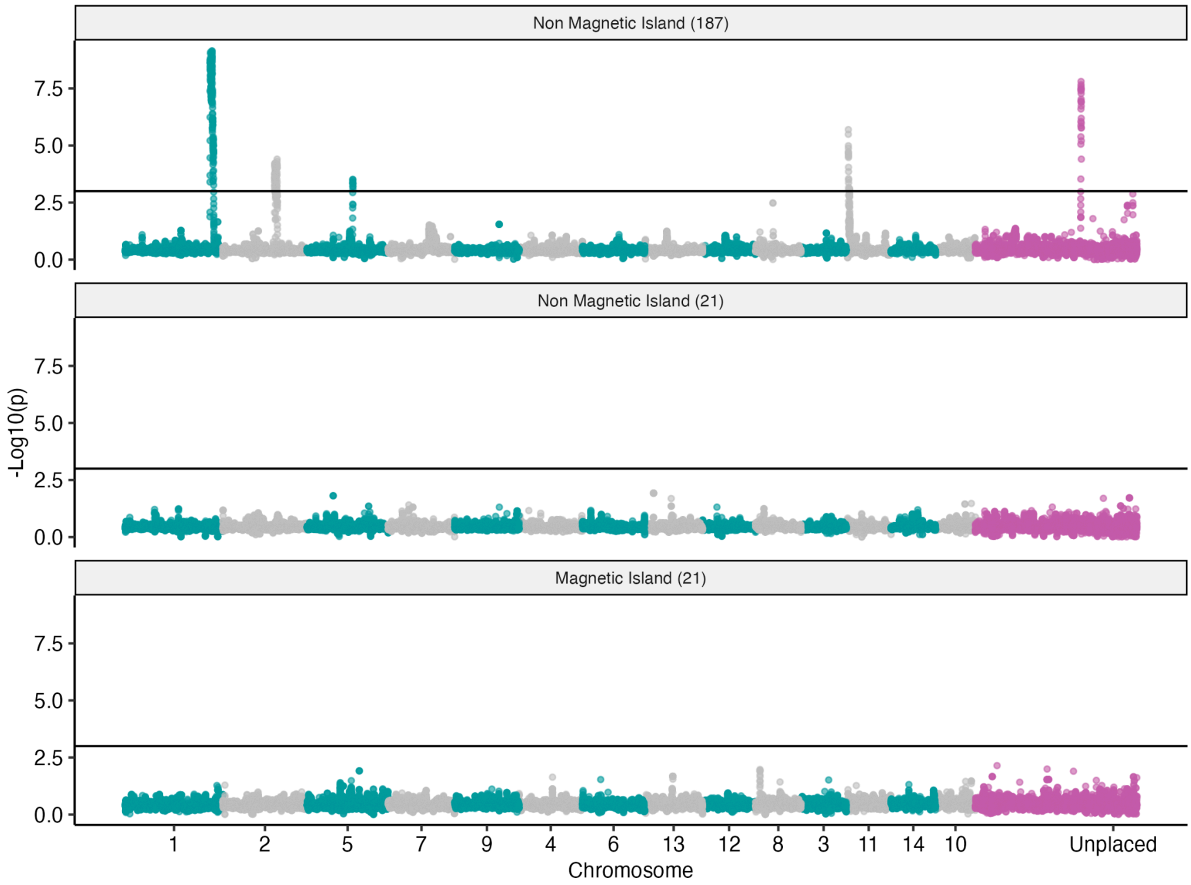


**Supplementary Figure S11:** Manhattan plots showing signals of local genetic structure in the *A. kenti* genome. Horizontal line in each plot shows a p=0.001 significance threshold. Top plot shows the results for all non-hybrid individuals in the non-Magnetic Island population, for which five highly significant peaks are present. Bottom two plots show a lack of significance at Magnetic Island (21 samples) and a random subset of 21 individuals from the non-Magnetic Island population.

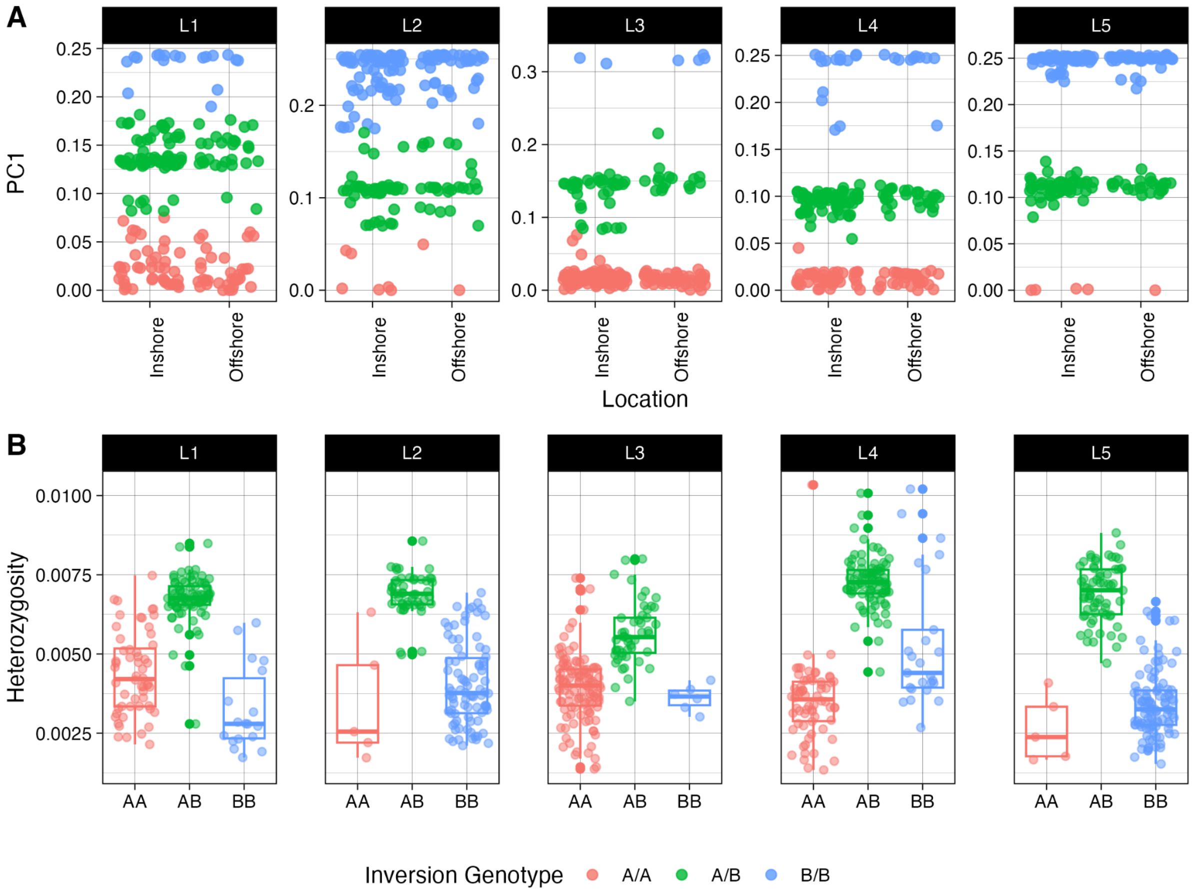


**Supplementary Figure S12:** Local genetic structure, genotype inference and heterozygosity within inversion loci. (A) Shows position of inshore and offshore samples along the first principal component (PC1) calculated with PCAngsd. (B) Shows individual heterozygosity for samples classified as AA, AB or BB for each inversion. Colors denote inferred genotypes for the inversion based on k-means clustering with K=3.

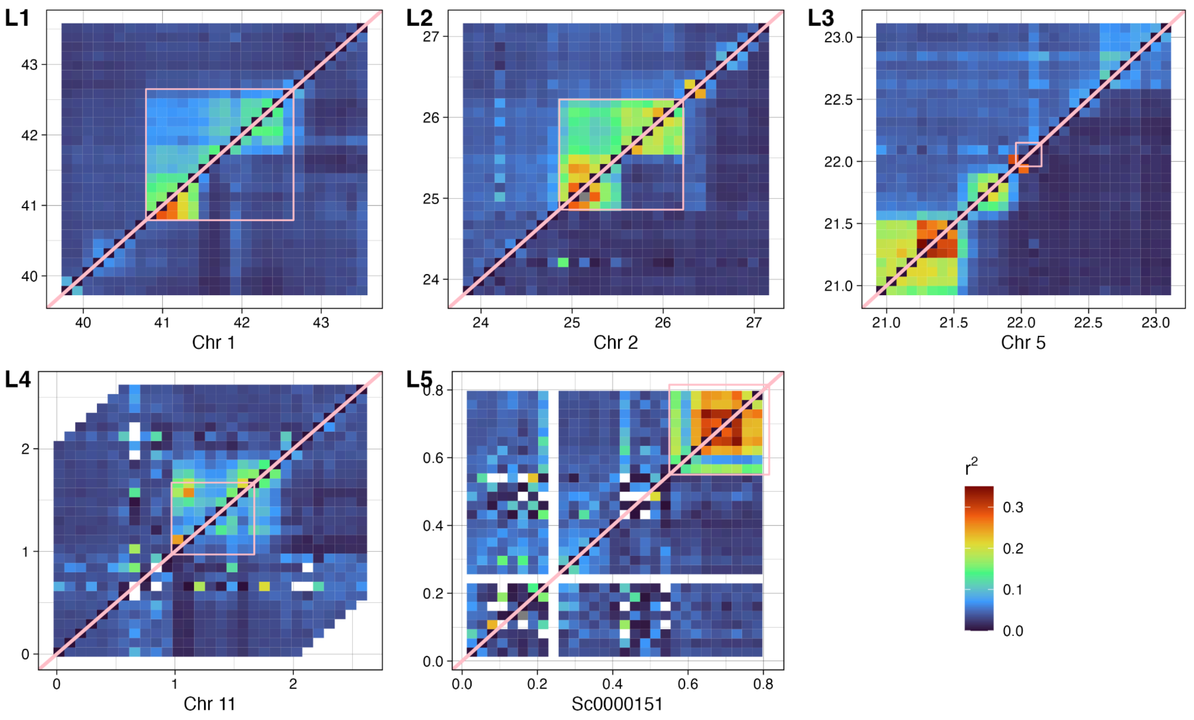


**Supplementary Figure S13:** Linkage disequilibrium at inversion loci calculated with ngsLD. Values in each pixel represent the average for all SNPs within the interval. All grid coordinates are based on pseudo-chromosome coordinates calculated with RagTag and are in units of megabases. Pink diagonal lines divide each plot into LD for homozygotes of the most common haplotype (bottom right) and heterozygotes (top left). Pink boxes delineate inversion boundaries as inferred from local population structure.

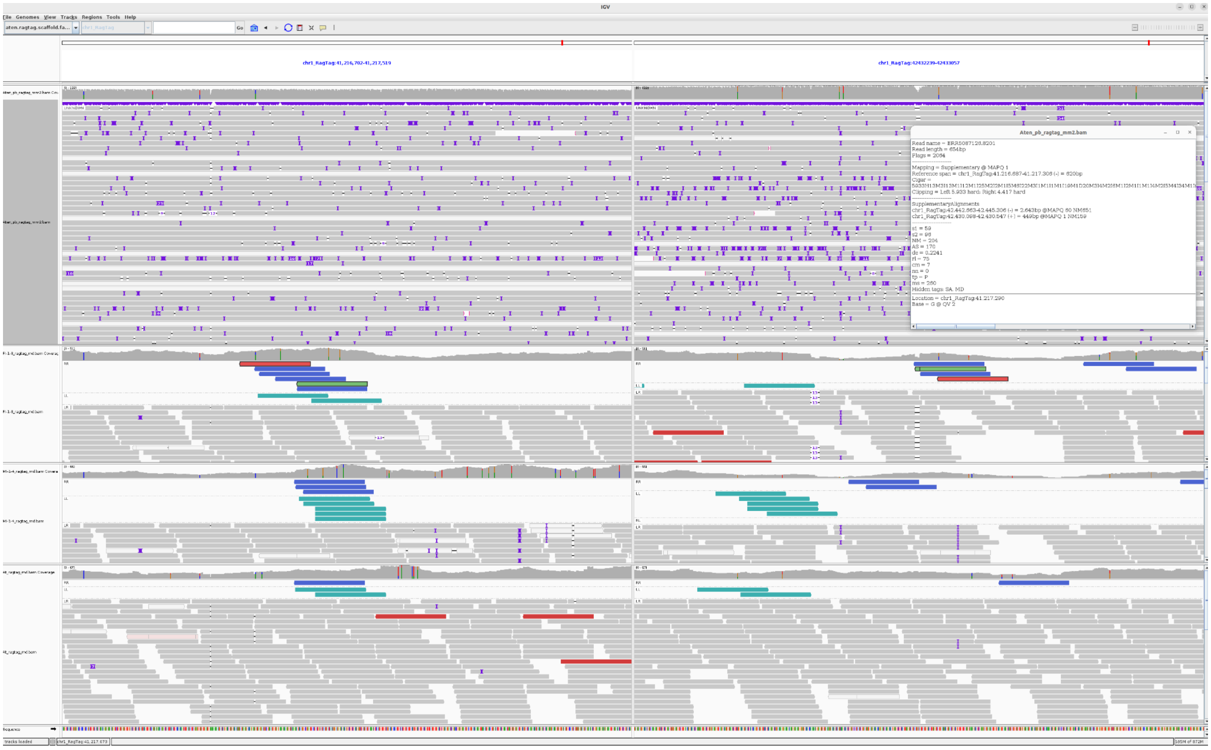

**Supplementary Figure S14:** Manual verification of the L1 inversion shows support in both short and long read sequence data with both breakpoints being captured. The top panel shows PacBio read alignments and a read split across the breakpoint highlighted by a pop up window showing the supplementary alignment across the breakpoint.  The bottom three panels show three coral genomes sequenced with short reads.  For the short reads, the reads pairs with orientations indicative of duplication breakpoints (RR and LL) are highlighted in blue and green.
